# The limited capacity of bioaerosols to serve as cloud-condensation nuclei may restrict their potential to initiate ice formation in mixed-phase clouds

**DOI:** 10.1101/2024.10.30.621021

**Authors:** Tina Šantl-Temkiv, Miha Živec, Marie Braad Lund, Mojca Benčina, Anja Bille Bohn, Samo Stanič, Griša Močnik

## Abstract

Bioaerosols are gaining prominence due to their ability to nucleate ice in clouds at high subzero temperatures, thereby impacting cloud characteristics, and longevity. Acting first as cloud condensation nuclei (CCN) and second as ice nucleating particles, bioaerosols induce ice formation through immersion freezing. Nevertheless, insights into bioaerosol ability to act as CCN *in situ* are currently lacking. We simultaneously collected bioaerosols from the condensed and interstitial phase of clouds using size selective inlets during autumn 2020 from the Otlica observatory (Slovenia). With a polarization Raman Lidar, we confirmed the coexistence of ice particles and liquid droplets during one of the three sampling campaigns. Using spectral flow cytometers, the PINGUIN cold-stage setup and MiSeq-facilitated amplicon sequencing to assess bacterial and fungal communities, we found that both bioaerosols and biogenic ice-nucleating particles were significantly more abundant in the interstitial phase compared to the condensed phase, suggesting that they were poor CCN, unable to compete with more hygroscopic particles. Most taxa did not exhibit preferential partitioning, suggesting that surface properties generally did not significantly influence their behaviour as CCN. Our results underline the need to systematically investigate the relation between hygroscopicity and ice-nucleating activity of bioaerosols to understand their *in-situ* effects on cloud formation.

**Synopsis:** The relationship between the ability of bioaerosols to nucleate cloud droplets and ice particles has not been determined. This study found that bioaerosols are poor cloud condensation nuclei which may impair their ability to nucleate ice and affect cloud formation and climate.

## 1. Introduction

Aerosol particles significantly impact cloud formation, properties, and lifetime, thus influencing the global climate ^1^. A subset of the aerosol particles, known as cloud condensation nuclei (CCN), are essential for forming cloud droplets in warm clouds, which develop at temperatures >0°C, and mixed-phase clouds, which develop at temperatures between 0°C and -38°C. The activity of CCN depends on their size and hygroscopicity, i.e. ability to attract water vapour ^2^. Another type of aerosol particles, called ice-nucleating particles (INP) are key players in the formation of ice in mixed-phase clouds. While CCN are common in the atmosphere (10^4^-10^6^ L^-1^ of air) ^2^, INP are much less abundant (10^-6^-10^3^ L^-1^ of air) ^3^. The type and concentration of CCN and INP directly reflect in cloud droplet number and size distribution, and therefore determine the optical properties of clouds which in-turn impacts the Earth’s radiation budget. Additionally, INP are crucial for much of continental precipitation^4^, influencing cloud lifetime and the atmospheric water cycle^5^. Despite the recognized key role of aerosols in controlling climate and the global water cycle, the response of cloud properties to the concentration and type of CCN and INP remains the major uncertainty when it comes to modelling climate change ^6^.

During the past decade, biological particles have received increasing attention due to their likely involvement in cloud development and precipitation ^7^. Laboratory studies have shown that, while mineral aerosols trigger ice nucleation at temperatures < -15 °C, bioaerosols nucleate ice at temperatures between -13°C and 0°C ^3^. These biological INP (bioINP) are primarily produced by ice-nucleation-active (INA) microorganisms, in particular epiphytic and soil bacterial strains affiliated to the genera *Pseudomonas*, *Erwinia*, *Pantoea*, *Xanthomonas*, and *Lysinibacillus* and soil fungal strains affiliated to the genera *Fusarium,* and *Mortierella*^7^. The majority of INA microorganisms produce highly potent INA proteins, which can either be aerosolized as parts of microbial cells ^8^ or disassociated from cells and bound to soil particles ^8–13^. In addition, bioINP of unknown origin, but with characteristics similar to INA proteins, were detected in the atmosphere and in precipitation^14^. While lidar and radar measurements show that ice is formed in ∼90% of clouds at temperatures above – 12°C ^15^, which is likely induced by bioINP, modelling studies are struggling to demonstrate the global and regional role of bioaerosols on cloud formation ^16–26^. To improve the ability of models to reproduce *in situ* measurements of ice formation, there is a need to improve our understanding of the quantitative impacts that bioINP play in mixed-phase clouds^27^.

The ability of bioaerosols to induce ice nucleation can be tightly linked to their ability to act as CCN. Field measurement- and modelling studies demonstrated that immersion freezing, where liquid water droplets must form before ice can nucleate, represents a predominant mode of ice nucleation^28^. Although few studies have examined the hygroscopicity of microbial cells as well as their ability to act as CCN, they are generally considered efficient cloud condensation nuclei due to their large size^27^, typically being categorized as large (0.2–2 μm) and giant (>2 μm) CCN^2^. Studies of single aerosolized microbial cells have shown that these have limited hygroscopicity^29,30^, which can however be greatly enhanced by the presence of salts on cell surfaces^30^. Cells of pure bacterial cultures were demonstrated to act as CCN at realistic supersaturations (0.07-1%)^31,32^. While Franc and DeMott (2002) found that only 25-30% of *Erwinia carotovora* cells activated at 1% supersaturation, Bauer et al. (2003) found that nearly all microorganisms that were isolated from atmospheric samples activated at much lower supersaturations (0.07- 0.11%). Surface properties of bacterial cells, such as hydrophobicity and charge, can be highly variable, and therefore these properties have been proposed to affect the ability of cells to activate into cloud droplets. While the existent studies have investigated defined bioaerosols at laboratory conditions^29–32^, competition with more hygroscopic aerosols for the available water vapour *in situ* may further complicate the ability of bioaerosols to act as CCN in clouds. Thus, modelling of mixed phase clouds, Simpson et al. (2018) demonstrated that INP were inefficient in competing for water vapor with more hygroscopic CCN, which ultimately limited INP in their immersion freezing potential^33^. The ability of bioaerosols to act as CCN and INP may therefore be linked tighter than what has previously been thought and it remains unclear how these properties synergistically affect ice-nucleation activity of bioaerosols *in situ*.

Our aim was to improve understanding of bioaerosols in terms of their ability to act as CCN and INP *in situ*. We thus combined lidar and ice-nucleation measurements with single-cell and next-generation sequencing techniques on samples of condensed and interstitial phase of clouds to test the following hypotheses: (i) the majority of bioaerosols, including bioINP, act as CCN and therefore partition into the condensed phase in liquid and mixed-phase clouds and (ii) the presence of bioINP in clouds coincides with the presence of ice particles in mixed-phase clouds at high subzero temperatures.

## 2. Methods

2.1. **The study location**

The campaign was conducted in the Vipava valley in western Slovenia, which is located on a junction between three different geographic regions with distinct climates. The valley is enclosed by the Vipava hills and the Karst plateau to the south, and the Trnovski gozd plateau and the Nanos to the north. The valley floor is level, ranging from approximately 60 m a.s.l. at its lowest to 170 m a.s.l. at its highest part. The northern edge ascends sharply to over 1000 m a.s.l., while the southern edge reaches around 300 m a.s.l. The terrain and infrastructure within and above the valley provide an ideal setting for studying aerosols in clouds, benefiting from seasonal stable and calm wind conditions. The clouds were sampled at the Atmospheric observatory of the University of Nova Gorica (UNG) at Otlica (45.93N, 13.91E, elevation 965 m a.s.l.) and were remotely probed by the UNG two-wavelength polarization Raman lidar located in Ajdovščina (45.89N, 13.91E), a town in the centre of the valley. The horizontal distance between the UNG lidar site and the Atmospheric observatory is approximately 5 km and a low cloud that can be observed remotely by lidar from Ajdovščina is the same cloud as the one sampled *in situ* at Otlica, allowing for simpler and more prolonged sampling periods compared to airborne missions.

### 2.2. Lidar measurements

Lidar is a versatile remote sensing tool exploiting backscattering of light pulses on atmospheric aerosols and molecules. Lidar has proven very useful in different atmospheric monitoring applications^34^. A polarization lidar, which emits coherent linearly polarized light and separates the return signal in parallel and cross-polarization states, observes the change in polarization of light due to different shapes of aerosols^35^. If the particles are spherical and homogeneous, the polarization of the backscattered light does not change, while nonspherical particles introduce a depolarized component into the backscattering signal^36^, allowing us to identify thermodynamic phases of water in clouds. The system employs two high-power Nd:YAG pulsed lasers at 355 nm and 1064 nm as transmitters and a 60 cm f/8 Cassegrain telescope as a receiver. The backscattered light collected with the telescope is split into five channels, each with a respective wavelength and polarization by a custom made polychromator. The lidar records the Mie–Rayleigh return signal at 1064 nm, the vibrational N_2_ Raman signal at 386.7 nm, the vibrational H_2_O Raman signal at 408.4 nm, and two Mie– Rayleigh signals at 355nm with different polarization planes. From the collected data, we obtained the extinction coefficient, backscatter coefficients, depolarization ratio, backscatter and extinction Ångström exponent, and lidar ratio profiles ^37^. To distinguish between liquid and solid phases of water in a cloud, the depolarization ratio of the 355 nm Mie–Rayleigh signals was used. Lidar measurements were performed simultaneously with aerosol sampling during three campaigns in October and December 2020. Aerosol loading and the depolarization ratio at the height of the observatory was obtained from the lidar data.

### 2.3. Sampling of bioaerosols in condensed and interstitial cloud phase

To investigate partitioning of bioaerosol particles between the condensed and the interstitial cloud phases during three sampling campaigns between Oct and Dec 2020 24 aerosol samples were collected (Table 1). Aerosols from the interstitial and the condensed phase were obtained simultaneously ∼2 m above ground (SI-Figure 1). The interstitial inlet was a large, inverted metal cylindrical container with a diameter of 63.6 cm and length 125 cm, closed at the top, save for the outlet connected to a BioSampler (SKC Ltd, flow rate of 11-12 lpm). The inlet was designed in such a way that the flow in it would be laminar and that the particles with a diameter larger than 5 μm (assuming a density of water droplets, corresponding both to most bioaerosols as well as cloud droplets), would settle rather than be sampled into the BioSampler (SI-Figure 2). After activating into cloud droplets, the diameters of CCN typically grow to 5–20 μm^38^ and would therefore not be sampled with this inlet due to settling. A virtual impactor (VI) was used to enhance the collection of supermicron particles, i.e. cloud droplets. The VI that was previously described by Drinovec et al. (2020) ^39^, was coupled to a BioSampler (SKC Ltd, flow rate of 11-12 lpm) and was operated at total-to-minor flow ratio of 6.3-7.3. SI-Figure 3 shows its concentration efficiency at this total-to-minor-flow ratio during testing with indoor aerosols of relevant diameters. The tubing connecting the inlets to the BioSamplers was sterilized with 70% ethanol prior to each of the three sampling campaigns. The sampling liquid in the BioSamplers was either a 0.9% (v:v) or 20% (v:v) NaCl solution (ambient temperatures <0°C, Table 1). The sampling liquid was autoclaved and filter sterilized (0.2 μm pore size), and thereafter split into 50 mL aliquots and stored frozen at -20°C until use. Prior to each sampling period, both BioSamplers were washed with MQ, sterilized by 70% ethanol and rinsed with sterile MQ. Twenty millilitres of sampling liquid were added to each BioSampler, which was then shaken to ensure contact with all inner surfaces. The resulting liquid was preserved as the sampling control. Immediately after sampling, we fixed samples for flow cytometry analysis with glutaraldehyde (final concentration of 2%) and stored them at -20°C. Samples used for the DNA and INP analysis were aliquoted and stored at -20°C.

**Figure 1:**
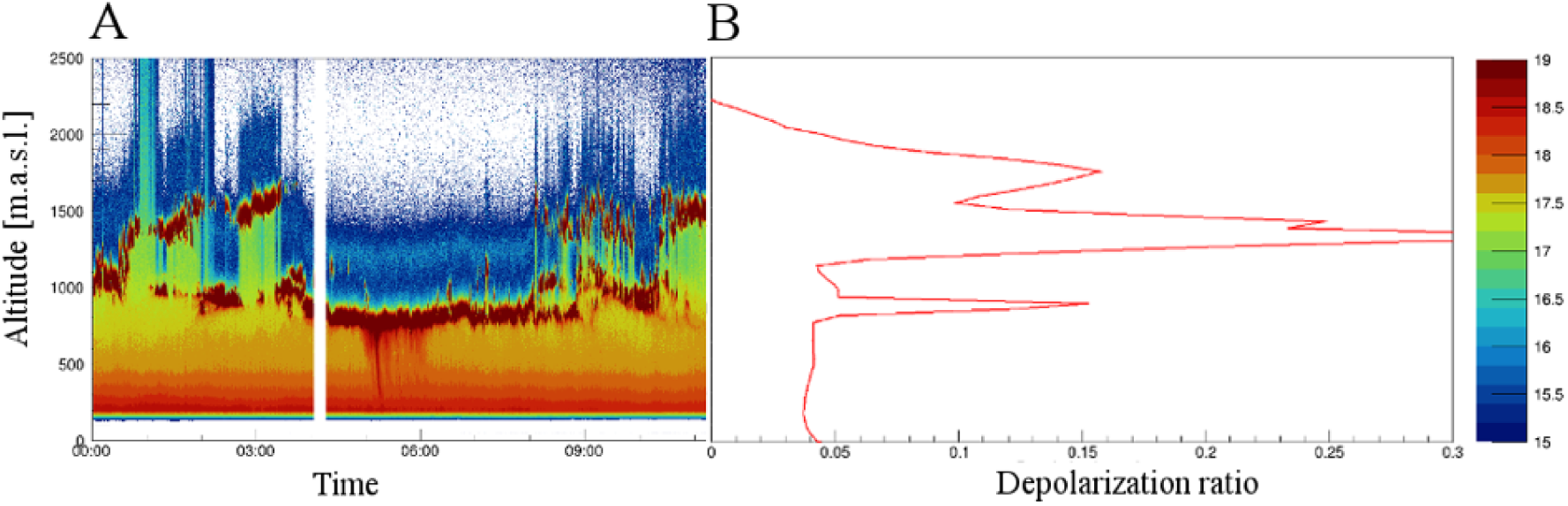
An example of the lidar analysis recorded on December 4^th^, 2020. (A) Log transformed data of the range corrected backscattered signal at 355 nm shows temporal variation of aerosol loading (arbitrary units) over the Vipava valley. In the morning, two layers of clouds were identified. The lower layer was at the same altitude as the Otlica observatory. (B) The depolarization ratio profile on December 4^th^ that was obtained between 1 am and 1:30 am also shows the presence of two cloud layers at 1000 m and 1500 m a.s.l. The depolarization of 15% in the lower layer at 1000 m altitude indicates that the clouds are mixed phase clouds. Alternatively, the higher depolarization ratio could result from multiple scattering of lidar pulses in a thick cloud layer. However, since the cloud layer was initially thin (≤200m), the number of multiply scattered photons should be low. This was confirmed by depolarization ratio values exceeding 30%, recorded in a thick elevated cloud layer (>1500 m a.s.l.).

**Figure 2:**
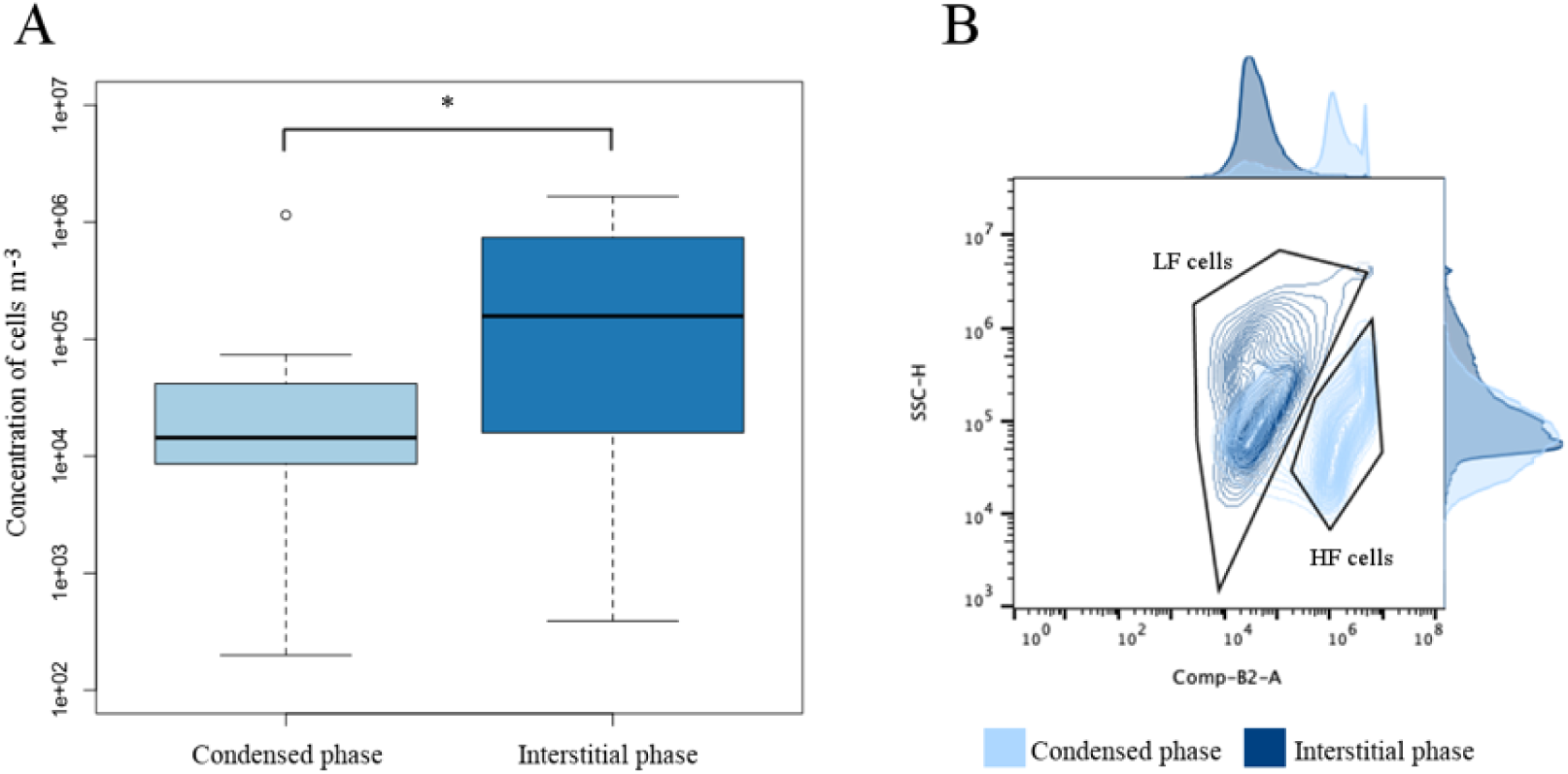
The concentration of microbial cells m^-3^ in the condensed and interstitial phase of the cloud as determined by flow cytometry. * indicates a significance level of <0.05. A contour plot obtained with the Aurora (Cytek Biosciences) spectral flow cytometer showing the side scatter of cells as a function of fluorescence in the B2 channel (emission wavelength of 528 nm, when excited with the blue laser) for a pair of condensed and interstitial samples (collected on Dec 19^th^, 2020). While LF (low fluorescent) cells are present in both phases, HF (high fluorescent) cells are present only in the condensed phase. Adjunct histograms are shown for each of the phases and parameters (side scatter and fluorescence in the B2 detector).

**Figure 3:**
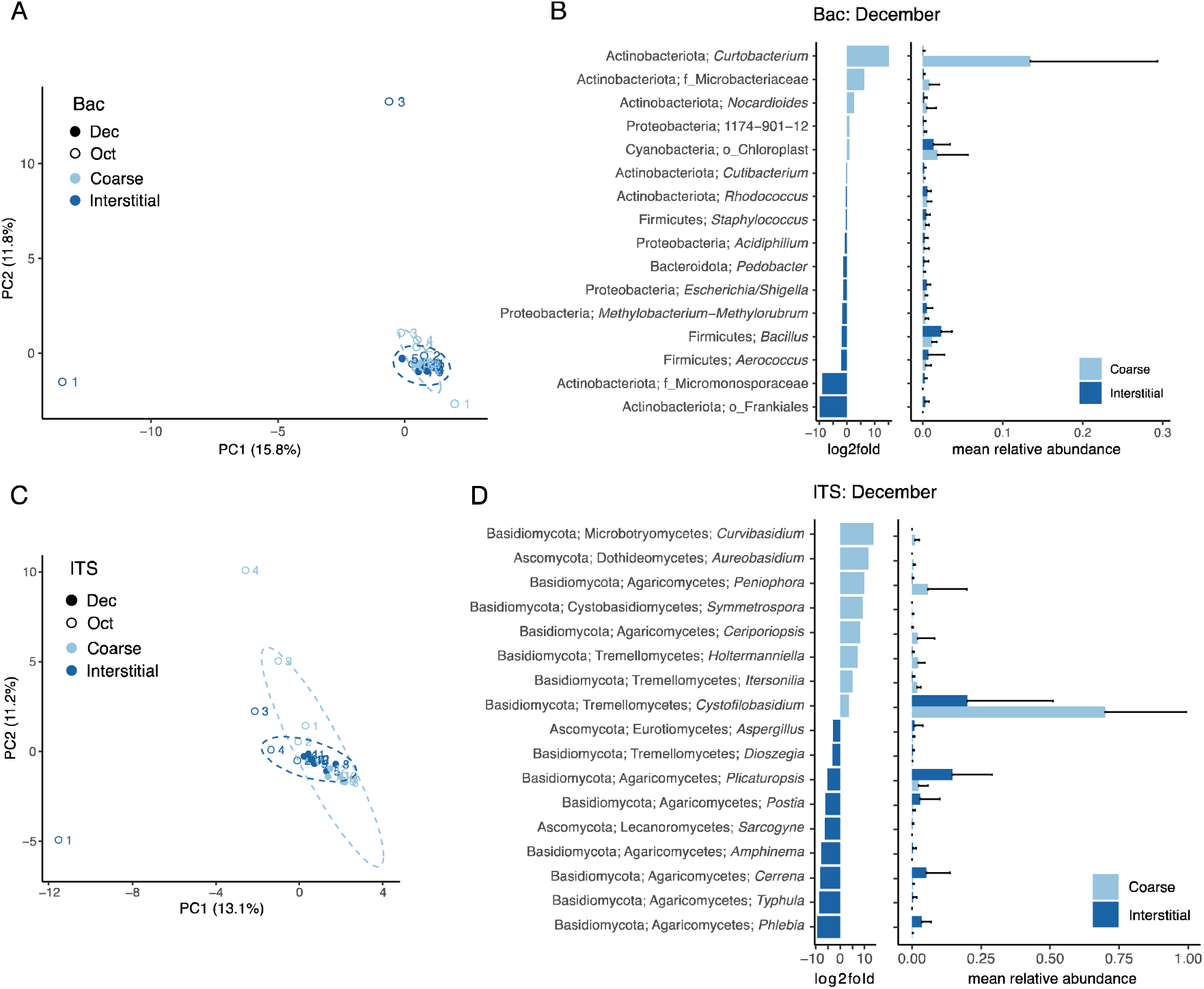
Analysis of bacterial 16S rRNA gene sequences (A, B) and fungal ITS sequences (C, D). (A, C) PCA plot of clr transformed read counts. Closed and open symbols show December and October samples, respectively. Colour coding shows condensed and interstitial samples according to legend. (B, D) Differentially abundant genera in December between condensed and interstitial samples (Wilcoxon signed rank tests, p<0.05). First bar plot shows log2-fold chance, second bar plot shows mean relative abundance (+SD).

**Table 1:**
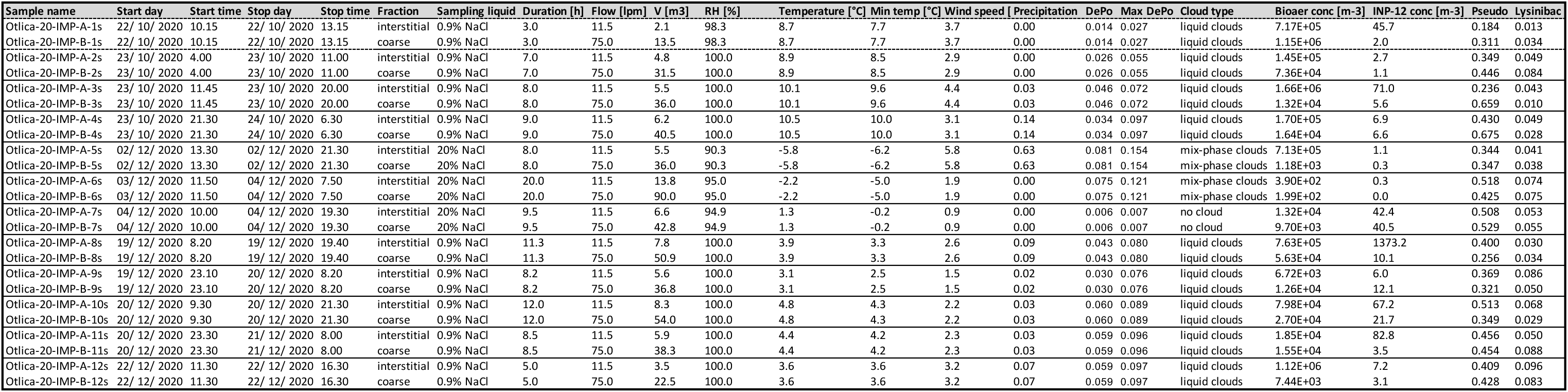
List of samples with the corresponding information on the sampling time, sampling liquid used in the impingers, sampling duration and flow, total volume of air sampled, meteorological conditions, lidar results, bioaerosol and INP-12 concentrations, and fraction of *Pseudomonas* and *Lysinibacillus* in the bacterial community.

### 2.4. Bioaerosol quantification and characterization of autofluorescence

The concentration and autofluorescence of microbial cells (which we term “bioaerosol concentration” throughout the article for simplification) were analysed by flow cytometry using 3-laser Aurora (Cytek Biosciences) and 5-laser ID7000 (Sony Biotechnology) spectral flow cytometers. A low threshold was set to remove debris from the analysis. Samples were stained with SYBR® Green I nucleic acid gel stain (10,000×, Sigma Aldrich) at a 5× final concentration. Stained samples were incubated at room temperature for 15 min in the dark prior to analysis. Unstained samples were used as controls and to analyse auto-fluorescent populations of particles in selected samples. We used SpectroFlo®, ID7000 ID_A_ and FlowJo (V10.8.1, Ch 2, Flowjo, LLC 2013-2016) software for data analysis and figure preparation.

### 2.5. DNA extraction amplicon sequencing

DNA was extracted from 250 μl aliquots of the individual samples using the DNeasy PowerSoil Pro Kit (Qiagen, Hilden, DE) following the protocol supplied by the manufacturer. The performance of the DNeasy PowerSoil Pro Kit has previously been demonstrated for atmospheric samples ^40^. The hypervariable regions V3 and V4 of the 16S rRNA gene and the fungal internal transcribed spacer (ITS) region were amplified using following the Illumina Metagenomic Sequencing Library Preparation guideline with slight modifications (see supplementary methods for details). For bacterial 16S rRNA genes the primers Bac341F (5’- CCT ACG GGN GGC WGC AG-3’) and Bac805R (5’-GAC TAC HVG GGT ATC TAA TCC-3’) ^41^ were used and for fungal ITS the primers ITS3_KYO2F (5’-GAT GAA GAA CGY AGY RAA-3’) and ITS4_KYO3R (5’-CTB TTV CCK CTT CAC TCG-3’) ^42^ were used. All amplicons were sequences on an Illumina MiSeq desktop sequencer using the V3 sequencing kit (Illumina) for 2x300 bp paired end sequencing.

The PCR mixture contained 20 μl template DNA, 2×KAPA HiFi Hotstart polymerase (KAPA Biosystems, Wilmington, MA), 0.2 μM forward primer, 0.2 μM reverse primer, and BSA (4 g/l). The thermal cycling consisted of an initial denaturation step at 95°C for 3 min, 30 cycles of denaturation at 95°C for 30 s, annealing at 55°C (16S rRNA) / 56 °C (ITS) for 30 s, elongation at 72 °C for 30 s and a final elongation step at 72 °C for 5 minutes. We used 30 μl AMPure XP magnetic beads to clean the PCR products. The second PCR was used to incorporate the Illumina overhang adaptors and was assembled in the same way only that 2 μl (16S rRNA) / 5 μl (ITS) of template was used in absence of BSA. The PCR was run for 10 cycles (16S rRNA) / 12 cycles (ITS) using the same conditions as for the first PCR. We used 20 μl AMPure XP magnetic beads to clean the PCR products. The third PCR was used to incorporate the Nextera XT Index primers. Each reaction contained 2.5 μl (16S rRNA) / 7.5 μl (ITS) of template, 2.5 μl of each index primers (N7XX and S5XX), 12.5 μl KAPA HiFi HotStart ReadyMix and dH2O up to the final volume of 25 μl. The PCR reaction was run for 9 cycles (16S rRNA) / 13 cycles (ITS) using the same conditions as for the first two PCR, after which the PCR product was cleaned with 56 μl AMPure XP beads. We used the Quant- iT™ dsDNA BR assay kit on a FLUOstar Omega fluorometric microplate reader (BMG LABTECH, Ortenberg, Germany) to quantify the PCR products. Then the products were diluted and pooled together in equimolar ratios. We quantified the pooled PCR products using the Quant-iT™ dsDNA BR assay kit on a Qubit fluorometer (Thermo Fisher Scientific, Waltham MA) and then sequenced them on the Illumina MiSeq platform (Illumina, San Diego, CA) using the Illumina V3 kit for paired end sequencing (2x300 bp).

### 2.6. Bioinformatic analysis

All analyses were done in R v 4.1.2 ^43^. Cutadapt v 0.1.1 ^44^ was used for trimming and primer removal. ‘DADA2’ v. 1.18.0 ^45^ was used for error correction, amplicon sequence variant (ASV) calling, chimera removal, and taxonomic classification. The Silva SSU reference database nr. 138 ^46^ was used for taxonomic classification of bacteria and the 2021 UNITE ‘general fasta release for fungi 2’ ^47^ was used for fungal ITS. Pooled sampling controls, nucleic acid extraction blanks and PCR negatives were used for decontaminating the data using the R package Decontam v 1.10.0 ^48^. Putative contaminants were identified using the prevalence method with a threshold of 0.1 and were subsequently removed from the data. All further data analysis were done using the R packages Phyloseq v. 1.34.0 ^49^, Microbiome v1.12.0 (Lahti), vegan v2.5.7 ^50^, ggplot2 v3.3.5 ^51^, dplyr ^52^, and custom R scripts.

Principal component analysis (PCA) and PERMANOVA was done on centred log-ratio (clr) transformed read counts (as implemented in the microbiome R package (Lahti)). Differential abundance analysis between interstitial and condensed samples were performed separately for October and December samples, using paired Wilcoxon signed rank test on clr transformed read counts. The p-values were adjusted for false discovery rate using the Benjamini & Hochberg method and 0.05 were used as a significance cutoff.

Sequence data has been deposited to GenBank under BioProject ID PRJNA1149942.

### 2.7. Cold-stage measurements

Droplet freezing experiments were performed over a temperature range between 0°C and - 28°C using the PINGUIN setup ^53^ to quantify the ice nucleating particles in the condensed and the interstitial samples. Either 48 (sampling controls) or 64 (samples) droplets of 30 µl were prepared in 384-well PCR plates, which was placed in the gallium bath cooled by two Peltier elements. Starting at ambient temperature, the temperature was decreased at 1 K min^-1^ until -28°C. Individual well temperatures were recorded using The FLIR A655sc w/25 thermal camera calibrated with a thermistor (TE Connectivity 2.252 kΩ) mounted in a fixed-point cavity. The freezing events were determined using a custom-made software based on the lateral heat release caused by the nucleation. The negative control for each run was MQ water filter-sterilized using a syringe filter (0.22-µm-pore size, Milipore, Massachusetts, USA). Sampling control for each sample was also included in the analysis. Based on the frozen fractions, we calculated the number of ice nucleation active sites per volume of air n (T) as proposed by ^54^: 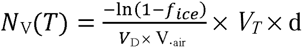. Where V_D_ the droplet volume, V_T_ the total volume of sample, f_ice_(T) the fraction of frozen droplets at a given temperature, d is the dilution factor and the V_air_ the volume of air collected for each sample. The freezing point depression caused by the NaCl present in the sampling liquid was calculated using the following equation: ΔT_f_ = n × c × E_f_ ^55^ . Where n is number of ions, c is molality of the salt and Ef is cryoscopic constant of water.

## 3. Results and discussion

### 3.1. Remote sensing of cloud ice

Based on the depolarization ratio retrieved directly from the lidar data (Figure 1), we analysed the properties of clouds during three measuring campaigns (22-24 October 2020, 1- 4 December 2020, and 19-22 December 2020, Table 1). During the first campaign (22-24 October 2020), the depolarization ratio of clouds at observatory height was low, with the maximum value of 5.5% (Table 1). Based on these values we conclude that the clouds were primarily composed of water droplets^56^. This was in line with average temperatures recorded in the clouds, which were between 8.7°C and 10.5°C and are consequently too high for ice crystal to form and sustain (Table 1). The RH recorded was on average 98.3% - 100.0%. During the second and third sampling campaign, the maximum depolarization ratio values in sampled clouds were higher, i.e. 12-16% (1-4 December 2020) and 7.5-10% (19-22 December 2020). During the second campaign, the average temperatures recorded in the clouds were between -5.8 and -2.2°C and the depolarization ratios confirmed the presence of nonspherical particles, which we interpret as ice particles. Since the depolarization ratios did not reach values typical of purely ice clouds, we conclude they were mixed phase clouds^56^. The presence of ice particles at high subzero temperatures would be best explained by the presence of biological INP (bioINP). The RH observed in the clouds during the second campaign was relatively low (on average 90.3%-95.0%), which may be linked either to the presence of dry air pockets, to a high ice water fraction, or to a combination of these two factors^57^. During the third campaign, the depolarization ratios were on the borderline between liquid and mixed-phase clouds. However, the average temperatures between 3.1 and 4.8 °C indicated predominantly liquid clouds. The average RH recorded was 100.0%.

### 3.2. The concentration of bioaerosols in interstitial and condensed phase

We determined bioaerosol concentration in the condensed and the interstitial phase by flow cytometry. The concentrations of bioaerosols were significantly (on average 30 times) higher in the samples than in the controls (Wilcoxon signed rank test, p-value <<0.001, SI-Figures 4 and 5). Thus, we found on average 4.5×10^5^ (min-max: 400-1.7×10^6^) bioaerosols m^-3^ in the interstitial phase and 1.2×10^5^ (min-max: 200-1.2×10^6^) bioaerosols m^-3^ in the condensed phase. These numbers are on the higher end of what has previously been found for bioaerosols across the globe ^58^ (SI-Figure 6, Table 1). There was a significant correlation between wind speed and total bioaerosol concentrations (Spearman’s rank correlation, p<0.01, N=12, SI-Figure 7A) and between temperature and bioaerosol concentrations in the condensed phase (Spearman’s rank correlation, p<0.0001, N=12, SI-Figure 7B). Wind speed and temperature have been linked to bioaerosol emissions, suggesting these bioaerosols may come from local sources ^62^.

**Figure 4:**
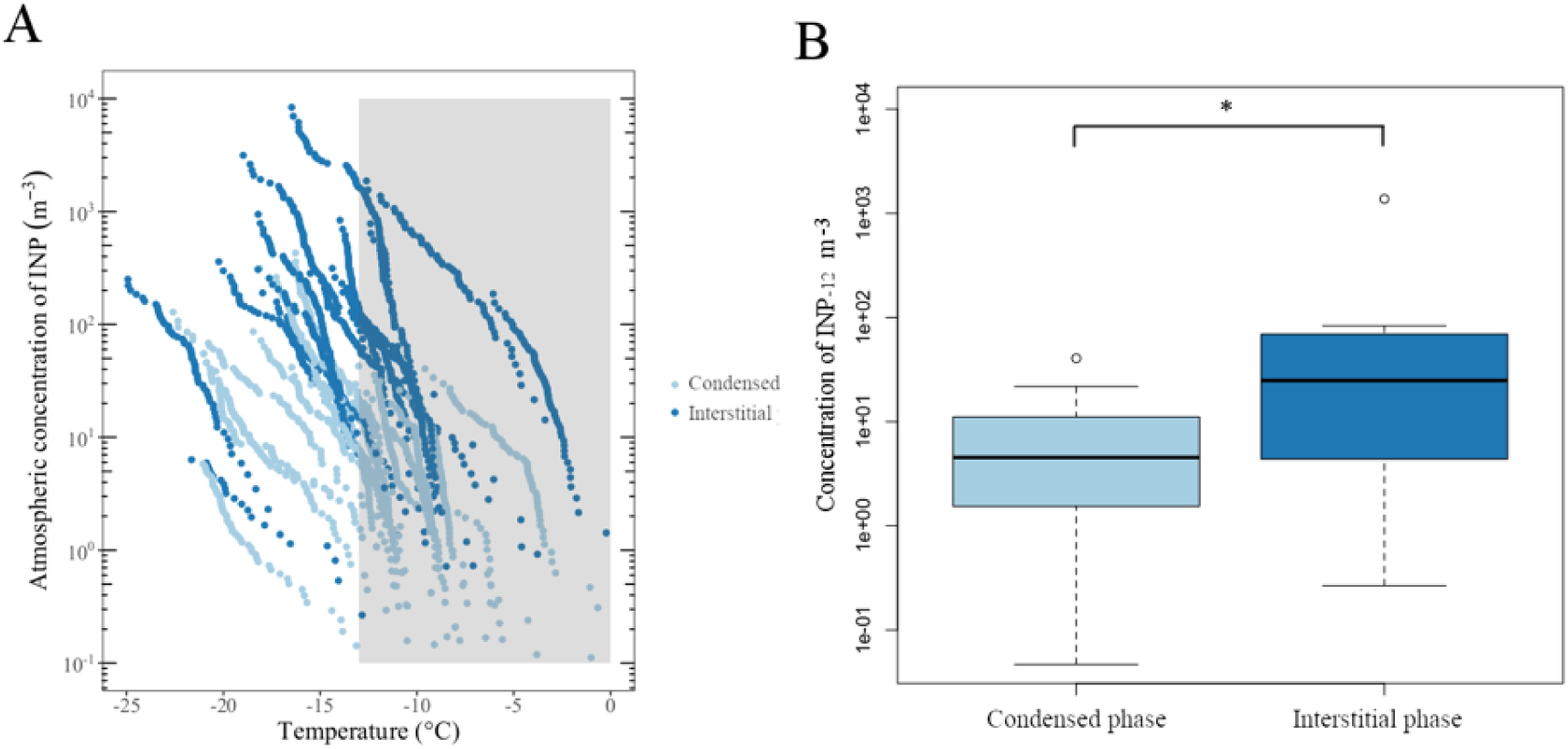
(A) Concentration of INP as a function of nucleation temperature in the condensed and the interstitial cloud phase. The shaded area denotes biogenic INP. (B) Concentration of INP_-12_ in the condensed and the interstitial cloud phase. * indicates a significance level of <0.05.

Bioaerosol concentrations in the interstitial phase were significantly higher (Wilcoxon signed rank test, p-value <0.05, on average 76 times higher, min-max: 0.6-602-times) than in the condensed phase (Figure 2A, SI-Figure 6), indicating that the majority of bioaerosols did not activate to cloud droplets at supersaturated conditions. Based on these results, we could not validate our first hypothesis and concluded that the majority of bioaerosols did not act as CCN and therefore did not partition into the condensed phase. This conclusion contradicts previous findings, claiming that large aerosols (>120 nm), are efficient CCN due to their large sizes ^69^. The size effect should apply to most airborne microbial cells, which are typically large compared to the majority of aerosols and have aerodynamic diameters >500 nm ^29,70^. It has previously been experimentally shown for selected microorganisms that they act as CCN at realistic supersaturations ^32^. Thus, Bauer et al. (2003) studied three microbial strains, which they isolated from aerosols and cloud water ^32^. They found that regardless of the strain, most cells (88-100%) activated already at low supersaturations (∼0.1%), demonstrating that some microbial cells are efficient CCN. Conversely, Franc and DeMott (2002) showed that four strains of *Erwinia carotovora*, isolated from plants and aerosols, exhibited significantly diminished ability to activate into droplets under realistic supersaturation conditions. Only 5-10% of the cells activated at 0.25% and 0.5% supersaturations, while 40-50% of ammonium sulphate particles activated at the same experimental conditions despite their much smaller sizes, which ranged between 10 and 200 nm ^31^. Variations in the ability of cells to act as CCN may be attributed to differences in their affinity to water, influenced by their surface properties ^71^. In line with this, Sharma and Rao (2002), measuring the contact angles between bacterial cells and water for 147 bacterial and yeast strains, demonstrated a large variability in hydrophilicity of microbial strains (contact angles of 16°-107°) ^72^. Unlike laboratory experiments that often involve only one type of aerosol, *in situ* conditions involve water vapor condensing on a diverse array of aerosols spanning a wide size range and exhibiting vastly different chemical properties. Under *in situ* conditions, aerosols are competing for water vapor while exposed to spatial and temporal heterogeneity with respect to the level of supersaturation ^33^. Despite the relatively high bioaerosol concentrations in the clouds that we studied, their concentration is still between 1 and 7 orders of magnitude smaller than the total concentration of CCN or concentration of cloud droplets ^2^. Typically, small CCN (60–200 nm), which are the most numerous, determine the concentration of droplets in a cloud. Thus, bioaerosols may not be able to compete successfully for the available water vapor with CCN that are more hygroscopic and much more numerous.

Spectral flow cytometry measurements of unstained samples revealed multiple populations of microbial cells based on autofluorescence (e.g. Figure 2B). Among these, we identified a major population distinct from the rest of the cells by its elevated fluorescence at 528 nm (high-fluorescent (HF) cells in Figure 2B, SI-Figure 8) and consistently present in the condensed phase collected during the second and third measurement campaigns. The concentration of HF cells was significantly higher in the condensed phase compared to the interstitial phase (Wilcoxon signed rank test, p < 0.05, SI-Figure 9), indicating a strong potential for HF cells to act as cloud condensation nuclei. When stained with Sybr Green and excited with a blue laser (488 nm), HF cells exhibited 8.3-14.8 times higher fluorescence at 528 nm compared to the regular population. They constituted 15% to 77% of bioaerosols in the condensed phase, as determined by the Aurora (Cytek Biosciences) spectral flow cytometer for stained samples. Analysing unstained samples using the ID7000 (Sony Biotechnology) spectral flow cytometer, we confirmed that the high fluorescence was due to autofluorescence, probably caused by the presence of autofluorescent biomolecules. The fluorescence of compounds, such as β-carotene, melanin, and flavin, corresponds well to what we observed in airborne cells (SI-Figure 8C) ^59–61^. A large fraction of airborne microbial cells have been shown to produce these types of pigments ^62–64^. In addition, we showed that HF cells exhibit a higher forward and side scatter compared to the other cells (SI-Figure 8B). As forward scatter typically correlates to cell size^65^, this indicates that the HF cells were either larger or formed aggregates which may mean that they act as giant CCN, improving their ability to activate into cloud droplets. Side scatted is instead linked to internal complexity of the cells, for example nucleus or granules^65^, and the high side scatter may indicate that the HF cells were not bacteria but were instead Eukaryotic cells. While overall most airborne microbial cells, which most likely had aerodynamic diameters >500 nm, were poor CCN, we found that only HF cells, which were bigger than most other microbial cells we observed, consistently activated into cloud droplets.

We observed differences in the bioaerosols concentrations in the condensed phase during the three campaigns (SI-Figure 10). During the second campaign, when ice particles and water droplets coexisted, the concentrations of bioaerosols in the condensed phase were lower than during the first and third campaign (SI-Figure 10). This suggests an ongoing Wegener– Bergeron process, where water is transferred from cloud droplets containing bioaerosols to ice particles, and therefore previously activated bioaerosols would shift back to the interstitial phase. The RH observed in the clouds during the second campaign was the lowest (on average 90.3%-95.0%), which may have also resulted in droplet evaporation. In addition, some of the activated bioaerosols may have acted as INP and have precipitated out of the cloud, which has previously been shown by field measurements and laboratory studies ^66–68^. Both of these processes could be responsible for the decreased concentration of bioaerosols in the condensed phase.

### 3.3. Selective bioaerosol partitioning between the interstitial and the condensed cloud phase

We used amplicon sequencing of bacterial 16S rRNA genes and the fungal ITS region to determine the community composition of microorganisms in the interstitial and the condensed phase, specifically search for the presence of INA genera, and determine whether specific taxa were better CCN than others, preferentially partitioning into the condensed cloud phase.

The bacterial communities were dominated by Gammaproteobacteria (57.4 ± 12.2 %), Alphaproteobacteria (13.5 ± 3.7%), Actinobacteria (10.2 ± 11.5%), Bacilli (11.4 ± 4.1%) and Cyanobacteria including Chloroplasts (3.4 ± 5.2%) (SI-Figure 11), which is in line with what has previously been found in the atmosphere ^7^. Among the different known INA bacterial genera, we detected *Erwinia*, *Pantoea*, *Xanthomonas*, *Lysinibacillus*, and *Pseudomonas* ^7^.

While members of the first three genera were present in only a few samples and at a small fraction (<0.5%), *Lysinibacillus* was present across all samples and accounted for 5.1±2.6% of the population in the interstitial and for 5.4±2.3% in the condensed phase (Table 1). The largest fraction of bacterial community was affiliated to *Pseudomonas* sp., with 39.3±10.5% in the interstitial and 43.3±13.2% in the condensed phase (Table 1). The proportion of *Lysinibacillus*, and *Pseudomonas* 16S rRNA genes in the interstitial phase were positively correlated (Spearman’s rank correlation, rho=0.64, p<0.05, SI-Figure 12), indicating that they were emitted from the same source. This is plausible as both genera may be found in the same environments. Strains affiliated with *Lysinibacillus* are often isolated from soils ^73^ and strains affiliated to *Pseudomonas* are among core members of the phyllosphere and rhizosphere microbiomes ^74^. Finally, the proportion of ASVs affiliated to the genus *Pseudomonas* in the condensed phase was negatively correlated to the depolarization ratio measures with the lidar, which may indicate selective precipitation of INA *Pseudomonas* strains from the clouds when ice formed in clouds as indicated by high depolarization ratios (SI-Figure 13).

The fungal communities were dominated by Basidiomycota affiliated to Agaricomycetes (46±33%) and Tremellomycetes (32±40%) (SI-Figure 14). Airborne cells affiliated to both classes were previously found to dominate fungal bioaerosol communities ^75–77^. Cells belonging to Ascomycota were more evenly distributed between 6 classes (Eurotiomycetes (6.7±15%), Leotiomycetes (3.4±8%), Sordariomycetes (2.9±6%), Dothideomycetes (1.2±2%), Lecanoromycetes (1.2±3%) and Saccharomycetes (0.6±2%)) (SI-Figure 14), which is also in agreement with previous results ^75,76^. Of the known INA fungal genera, *Fusarium* was only detected in one of the samples, while *Mortierella* was not detected in any sample.

Since fungal INA proteins can dissociate from the cells and bind to soil particles^7^, the absence of INA fungal strains does not necessarily mean that fungal INA proteins are absent. Based on a PCA analysis, we concluded that the communities in the interstitial and the condensed phases are highly similar for both bacteria and fungi (Figure 3A and C).

Conversely, a PERMANOVA analysis showed no significant differences between interstitial and condensed communities for bacteria (p > 0.05), but a significant difference in the fungal communities (p < 0.05). This difference was mainly driven by few samples (Figure 3C).

Šantl-Temkiv et al. (2015) demonstrated that cells of identical strains affiliated to rain-borne *Pseudomonas* sp. changed their surfaces properties due to chemical and physical stress from hydrophilic (contact angles of 20°-60°) to hydrophobic (65°-100°) ^8^. Thus, differences in the cell’s surface properties due to their life history could explain why some cells of specific taxa, present in the air, were activated into cloud droplets while most cells of the same taxa remained in the interstitial phase.

To pinpoint taxa that are particularly good or bad CCN, we performed a differential abundance analysis. As we observed that the community composition differed, for fungi, between samples collected in October and December, we performed this analysis separately on communities collected during these two periods. In October, none of the bacterial or fungal genera were differentially abundant between the interstitial and condensed phase of the cloud. In December, there were 16 bacterial and 17 fungal genera (out of 379 and 236 in total, respectively) that were differentially abundant in one of the cloud phases (Figures 3B,D). Most of the differentially abundant bacterial genera represented a minor fraction of the total community (< 1.8% on average) except for the genus *Curtobacterium* that had a mean relative abundance of 13.5% (±15.9) and 0.1% (±0.2) in the condensed and interstitial phases, respectively. *Curtobacterium* is a cosmopolitan bacterial genus that is commonly found on plant surfaces and in soils, and that was previously also found in the atmosphere ^78^.

Like bacteria, most of the differentially abundant fungal genera represented a minor fraction of the community (<3% on average). Two genera had a higher mean relative abundance. *Plicaturopsis*, which had a mean relative abundance of 2.4% (±3.4) and 14.8% (±14.2) in the condensed and interstitial phases, respectively, and thus seems to be a poor CCN. *Cystofilobasidium* had a mean relative abundance of 69.9 % (±29.4) and 20.0% (±31.0) in the condensed and interstitial phases, respectively. Species affiliated to *Cystofilobasidium* primarily occur as yeast and are found in both marine and terrestrial environments^79^. Interestingly, analysing microorganisms associated with highly INA aspen leaf litter, Vaseby et al. (2019) detected *Curtobacterium* sp. together with the INA bacteria *Pseudomonas syringae*, and *Pantoea ananatis* as well as found that strains affiliated to *Cystofilobasidium* sp. dominated the fungal community^80^. This indicates a potential common source for the *Cystofilobasidium* sp., *Curtobacterium* sp. and the INA taxa in our samples. Both *Curtobacterium* sp. and *Cystofilobasidium* sp. were present in the same samples from the condensed phase as the HF cells. Therefore, we suggest that cells affiliated to these two genera constitute the HF cell population with an enhanced potential to act as CNN. Strains affiliating to *Curtobacterium* ^81,82^ and *Cystofilobasidium* sp. ^79,83^ were shown to synthesize carotenoid pigments that are photoprotective ^84^. However, as many microbial taxa in the atmosphere contain carotenoids ^62–64^ this is barely the sole explanation of the increased autofluorescence observed for the HF cells. Alternatively, our results could be explained by the presence of flavins, which have similar fluorescent properties as carotenes. Thus, Surre et al. (2018) demonstrated an evolutionary-conserved stress response in both bacteria and Eukaryotes that leads to increased autofluorescence caused by oxidation or synthesis of flavins that are involved in the detoxification of reactive oxygen species ^61^. A prolonged residence time in the atmosphere, which is an oxidative environment, may lead to the oxidation of flavins in airborne cells, resulting in increased autofluorescence in the HF cells. Prolonged atmospheric residence time may also affect the ability of bioaerosols to act as CCN, as the aging of atmospheric aerosols enhances their ability to act as CCN ^85^. This may explain why the HF cells were nearly exclusively found in the condensed phase of the clouds.

### 3.4. Partitioning of ice-nucleating particles between the interstitial and the condensed cloud phase

We assessed the concentration of INP as a function of their nucleation temperature in both the interstitial and the condensed phase of the mixed-phase clouds using the PINGUIN cold-stage instrument. The frozen fractions are shown in SI-Figure 15. The MQ controls and sample controls were clearly separated from the sample (SI-Figure 15). We found that most samples (83%) contained INP that were active at temperatures ≥–12°C (Figure 4A, SI-Figure 16), indicating that these were bioINP likely containing INA proteins. There was no significant correlation between temperature or wind speed and the concentration of INPs active at ≥–12°C (INP_-12_). However, the positive correlation between the INP_-12_ and the bioaerosol concentration (Spearman’s rank correlation, rho=0.4, p<0.05, SI-Figure 17) supports their biogenic origin. There was no significant correlation between INP_-12_ and the fraction of the predominant known INA bacterial genera, i.e. *Pseudomonas* and *Lysinobacillus*.

There were on average 142.2 INP_-12_ m^-3^ of air (min-man: 0.3-1373.2, Figure 4B, SI-Figure 15, Table 1) in the interstitial aerosols, which is at the higher end of previously reported INP_-12_ concentrations worldwide ^14^. High concentrations of INP_-12_ were in line with high concentrations of bioaerosols. Elevated concentrations of INP were previously recorded in clouds compared to outside the clouds ^86^ and positively correlated with high relative humidity ^87^. We observed on average 8.9 INP_-12_ m^-3^ of air (min-man: 0.05-40.5) in the condensed phase of clouds (Figure 4B, Table 1). These concentrations were at the lower end of the INP_-12_ concentrations found in cloud water ^88^ or estimated based on measurements in precipitation^89^. As with bioaerosols, we found that there were significantly lower concentrations of INP_-12_in the condensed than in the interstitial phase (Wilcoxon signed rank test, p-value <0.01). This suggests that the ability of bioINP to act as CCN was low, as we have also observed for bioaerosols, further underlining that we could not validate our first hypothesis. A modelling study previously established that INP were inefficient CCN when competing with very hygroscopic CCN for water vapor ^33^. As activation of INPs into cloud droplets is required to induce immersion freezing, which is a predominant mode of ice nucleation ^28^, lacking the ability to act as CCN could compromise their capacity to induce ice formation *in situ*.

Based on the lidar measurements, we conclude that we investigated liquid clouds during the first and third campaign when the temperatures were >0°C, while we investigated mixed-phase clouds during the second campaign when there were subzero temperatures. We therefore validated the second hypothesis stating that the presence of bioINP in clouds coincides with the presence of ice particles in mixed-phase clouds at high subzero temperatures. Alternatively, the external seeder–feeder process ^90^, i.e. ice crystals precipitating from a higher cloud layer (∼1500 m) into the lower cloud layer (∼1000 m), was responsible for the observed ice particles. However, considering the typical temperature lapse rates ^91^ of up to a maximum 6.5°C km^-1^ and potentially lower, the upper cloud layer would still be in a temperature range of >-13°C, where the formation of ice particles is driven by bioINP. As we observed precipitation (<1 mm) during the same campaign, this indicates that bioINP was also involved in triggering precipitation formation.

Finally, we observed differences in the INP_-12_ concentrations during the three campaigns (SI-Figure 18). During the second campaign when the presence of INP_-12_ coincided with the presence of ice particles in clouds, the concentrations of INP_-12_ in clouds were lower than during the first and third campaign (SI-Figure 18). This may indicate that INP were preferentially precipitated from the cloud as has previously been reported based on field measurements and laboratory studies ^66–68^. This was corroborated by the precipitation we observed during the second campaign. Lastly, the concentration of INP in the condensed phase of clouds decreased between the first and the second day of the second campaign, indicating that bioINP preferentially precipitated from clouds in rain.

## Summary and Conclusions

Using the depolarization ratio based on the lidar data, we concluded that we investigated liquid clouds during two of the measurement campaigns and mixed-phase clouds in one of the campaigns. We found a high concentration of bioaerosols and bioINP was characteristic for the clouds. Strikingly, we observed for the first time that both bioaerosols and bioINP were predominantly found in the interstitial phase of the clouds therefore suggesting they were poor CCN under the *in situ* conditions, where they likely competed with highly hygroscopic CCN. Poor ability to act as CCN reduces the capability of these bioINP to induce immersion freezing. Within the microbial community, we identified strains affiliating with known bacterial INA genera, i.e. *Pseudomonas* and *Lysinobacillus* which were present at high proportions of 41% and 5%, respectively, and which may be associated with the high concentrations of bioINP that we observed. We established the co-occurrence of bioINP, cloud ice particles, and precipitation at high subzero temperatures during one of the measurement campaigns, pointing to the fact that bioINP likely induced ice formation and precipitation. For most bacterial and fungal taxa, cells had comparable properties with respect to their potential to act as CCN. They were poor CCN regardless of their size, physiology or surface properties, as they were present in similar proportions in the interstitial and the condensed phase. Notable exceptions were cells of *Cystofilobasidium* sp. and *Curtobacterium* sp. that as good CCN were preferentially found in the condensed phase. These cells were larger in size which may be one of the reasons for their ability to act as good CCN. In addition, they were characterized by an elevated autofluorescence, likely caused by oxidized flavins. This indicates that atmospheric aging enhanced their ability to act as CCN. This is the first study that links autofluorescence to surface properties and CCN in bioaerosols, solicitating further studies of this association. Overall, we found that both bioaerosols and bioINP were poor CCN *in situ*, which could limit their ability to nucleate cloud ice. We conclude that there is a link between hygroscopicity and ice-nucleating activity of bioaerosols and that this link must be systematically investigated to better understand their *in-situ* effects on cloud formation.

## Author contributions

TŠT and GM designed and supervised the research project. TŠT collected the samples, conducted the flow cytometry analysis, and supervised the DNA extraction and sequencing. MŽ set up the lidar measurements and analysed the lidar data supervised by SS. MBL preformed the bioinformatic analysis. MB and ABB helped with measurements using the spectral flow cytometers. The manuscript was written by TŠT and revised by all co-authors.

## Acknowledgements

The authors are grateful to B. Poulsen and S. Nielsen for excellent technical assistance and to R. Krapež for his logistic support. Flow cytometry using Sony ID7000 was performed at the FACS Core Facility, Aarhus University, Denmark. We thank L. Wang for his help with lidar data analysis, and K. Finster for his constructive review of the manuscript. This research was co-funded by the Slovenian Research Agency (programs P1-0031, P1-0385 and I0-0033), the Public Scholarship, Development, Disability and Maintenance Fund of Republic of Slovenia (11013-21), The Danish National Research Foundation (DNRF106, to the Stellar Astrophysics Centre, Aarhus University), the Novo Nordisk Foundation (NNF19OC0056963), the Villum Fonden (23175 and 37435), and the Independent Research Fund Denmark (9145-00001B).

## References

1. Boucher, O.; Randall, D.; Artaxo, P.; Bretherton, C.; Feingold, G.; Forster, P.; Kerminen, V.-M.; Kondo, Y.; Liao, H.; Lohmann, U.; Rasch, P.; Satheesh, S. K.; Sherwood, S.; Stevens, B.; Zhang, X. Y. Clouds and Aerosols. In: Climate Change 2013: The Physical Science Basis. Contribution of Working Group I to the Fifth Assessment; Cambridge University Press, Cambridge, United Kingdom and New York, NY, USA, 2013; pp 571–658. 10.1017/CBO9781107415324.016.

(2) Pruppacher H.R.; Klett J. D. Microphysics of Clouds and Precipitation; 1980; Vol. 31. 10.1088/0031-9112/31/5/037.

(3) Kanji, Z. A.; Ladino, L. A.; Wex, H.; Boose, Y.; Burkert-Kohn, M.; Cziczo, D. J.; Krämer, M. Overview of Ice Nucleating Particles. Meteorol. Monogr. 2017, *58*, 1.1–1.33. 10.1175/AMSMONOGRAPHS-D-16-0006.1.

(4) Mülmenstädt, J.; Sourdeval, O.; Delanoë, J.; Quaas, J. Frequency of Occurrence of Rain from Liquid-, Mixed-, and Ice-Phase Clouds Derived from A-Train Satellite Retrievals. Geophys. Res. Lett. 2015, 42, 2015GL064604. 10.1002/2015gl064604.

(5) Morris, C. E.; Georgakopoulos, D. G.; Sands, D. C. Ice Nucleation Active Bacteria and Their Potential Role in Precipitation. J. Phys. IV Proc. 2004, 121, 87–103. 10.1051/jp4:2004121004.

(6) Short-Lived Climate Forcers. In Climate Change 2021 – The Physical Science Basis: Working Group I Contribution to the Sixth Assessment Report of the Intergovernmental Panel on Climate Change; Intergovernmental Panel on Climate Change (IPCC), Ed.; Cambridge University Press: Cambridge, 2023; pp 817–922. 10.1017/9781009157896.008.

(7) Šantl-Temkiv, T.; Amato, P.; Casamayor, E. O.; Lee, P. K. H.; Pointing. Microbial Ecology of the Atmosphere. FEMS Microbiol. Rev. 2022, *fuac009*. 10.1093/femsre/fuac009.

(8) Šantl-Temkiv, T.; Sahyoun, M.; Finster, K.; Hartmann, S.; Augustin-Bauditz, S.; Stratmann, F.; Wex, H.; Clauss, T.; Nielsen, N. W.; Sørensen, J. H.; Korsholm, U. S.; Wick, L. Y.; Karlson, U. G. Characterization of Airborne Ice-Nucleation-Active Bacteria and Bacterial Fragments. Atmos. Environ. 2015, 109, 105–117. 10.1016/j.atmosenv.2015.02.060.

(9) O’Sullivan, D.; Murray, B. J.; Ross, J. F.; Whale, T. F.; Price, H. C.; Atkinson, J. D.; Umo, N. S.; Webb, M. E. The Relevance of Nanoscale Biological Fragments for Ice Nucleation in Clouds. Sci. Rep. 2015, 5, 1–7. 10.1038/srep08082.

(10) Wilson, T. W.; Ladino, L. A.; Alpert, P. A.; Breckels, M. N.; Brooks, I. M.; Browse, J.; Burrows, S. M.; Carslaw, K. S.; Huffman, J. A.; Judd, C.; Kilthau, W. P.; Mason, R. H.; McFiggans, G.; Miller, L. A.; Najera, J. J.; Polishchuk, E.; Rae, S.; Schiller, C. L.; Si, M.; Temprado, J. V.; Whale, T. F.; Wong, J. P. S.; Wurl, O.; Yakobi-Hancock, J. D.; Abbatt, J. P. D.; Aller, J. Y.; Bertram, A. K.; Knopf, D. A.; Murray, B. J. A Marine Biogenic Source of Atmospheric Ice-Nucleating Particles. Nature 2015, 525 (7568), 234–238. 10.1038/nature14986.

(11) Conen, F.; Morris, C. E.; Leifeld, J.; Yakutin, M. V.; Alewell, C. Biological Residues Define the Ice Nucleation Properties of Soil Dust. *Atmospheric Chem*. Phys. 2011, 11 (18), 9643–9648. 10.5194/acp-11-9643-2011.

(12) Irish, V. E.; Elizondo, P.; Chen, J.; Choul, C.; Charette, J.; Lizotte, M.; Ladino, L. A.; Wilson, T. W.; Gosselin, M.; Murray, B. J.; Polishchuk, E.; Abbatt, J. P. D.; Miller, L. A.; Bertram, A. K. Ice-Nucleating Particles in Canadian Arctic Sea-Surface Microlayer and Bulk Seawater. *Atmospheric Chem*. Phys. 2017. 10.5194/acp-17-10583-2017.

(13) Creamean, J. M.; Kirpes, R. M.; Pratt, K. A.; Spada, N. J.; Maahn, M.; de Boer, G.; Schnell, R. C.; China, S. Marine and Terrestrial Influences on Ice Nucleating Particles during Continuous Springtime Measurements in an Arctic Oilfield Location. *Atmospheric Chem*. Phys. 2018, 18, 18023–18042. 10.5194/acp-2018-545.

(14) Kanji, Z. A.; Ladino, L. A.; Wex, H.; Boose, Y.; Burkert-Kohn, M.; Cziczo, D. J.; Krämer, M.; Kanji, Z. A.; Ladino, L. A.; Wex, H.; Boose, Y.; Burkert-Kohn, M.; Cziczo, D. J.; Krämer, M. Overview of Ice Nucleating Particles. Meteorol. Monogr. 2017, *58*, 1.1–1.33. 10.1175/AMSMONOGRAPHS-D-16-0006.1.

(15) Bühl, J.; Ansmann, A.; Seifert, P.; Baars, H.; Engelmann, R. Toward a Quantitative Characterization of Heterogeneous Ice Formation with Lidar/Radar: Comparison of CALIPSO/CloudSat with Ground-Based Observations. Geophys. Res. Lett. 2013, 40 (16), 4404–4408. 10.1002/grl.50792.

(16) Hoose, C.; Kristjánsson, J. E.; Burrows, S. M. How Important Is Biological Ice Nucleation in Clouds on a Global Scale? Environ. Res. Lett. 2010, 5 (2). 10.1088/1748-9326/5/2/024009.

(17) Hoose, C.; Kristjánsson, J. E.; Chen, J.-P.; Hazra, A. A Classical-Theory-Based Parameterization of Heterogeneous Ice Nucleation by Mineral Dust, Soot, and Biological Particles in a Global Climate Model. J. Atmospheric Sci. 2010, 67 (8), 2483– 2503. 10.1175/2010JAS3425.1.

(18) Sahyoun, M.; Wex, H.; Gosewinkel, U.; Šantl-Temkiv, T.; Nielsen, N. W.; Finster, K.; Sørensen, J. H.; Stratmann, F.; Korsholm, U. S. On the Usage of Classical Nucleation Theory in Quantification of the Impact of Bacterial INP on Weather and Climate. Atmos. Environ. 2016, 139, 230–240. 10.1016/j.atmosenv.2016.05.034.

(19) Sahyoun, M.; Korsholm, U. S.; Sørensen, J. H.; Šantl-Temkiv, T.; Finster, K.; Gosewinkel, U.; Nielsen, N. W. Impact of Bacterial Ice Nucleating Particles on Weather Predicted by a Numerical Weather Prediction Model. Atmos. Environ. 2017, 170. 10.1016/j.atmosenv.2017.09.029.

(20) Sesartic, A.; Lohmann, U.; Storelvmo, T. Bacteria in the ECHAM5-HAM Global Climate Model. Atmos Chem Phys 2012, 12 (18), 8645–8661. 10.5194/acp-12-8645-2012.

(21) Sesartic, A.; Lohmann, U.; Storelvmo, T. Modelling the Impact of Fungal Spore Ice Nuclei on Clouds and Precipitation. 2013, 014029. 10.1088/1748-9326/8/1/014029.

22. Sesartic, A.; Lohmann, U.; Storelvmo, T. Bacteria in the ECHAM5-HAM Global Climate Model. 2011. 10.5194/acpd-11-1457-2011.

23. Phillips, V. T. J.; Andronache, C.; Christner, B.; Morris, C. E.; Sands, D. C.; Bansemer, A.; Lauer, A. Potential Impacts from Biological Aerosols on Ensembles of Continental Clouds Simulated Numerically. 2009, 987–1014.

(24) Hummel, M.; Hoose, C.; Pummer, B.; Schaupp, C.; Fröhlich-Nowoisky, J.; Möhler, O. Simulating the Influence of Primary Biological Aerosol Particles on Clouds by Heterogeneous Ice Nucleation. *Atmospheric Chem*. Phys. 2018, 18, 15437–15450. 10.5194/acp-18-15437-2018.

(25) Spracklen, D. V.; Heald, C. L. The Contribution of Fungal Spores and Bacteria to Regional and Global Aerosol Number and Ice Nucleation Immersion Freezing Rates. *Atmospheric Chem*. Phys. 2014, 14 (17). 10.5194/acp-14-9051-2014.

(26) Vergara-Temprado, J.; Murray, B. J.; Wilson, T. W.; O’Sullivan, D.; Browse, J.; Pringle, K. J.; Ardon-Dryer, K.; Bertram, A. K.; Burrows, S. M.; Ceburnis, D.; Demott, P. J.; Mason, R. H.; O’Dowd, C. D.; Rinaldi, M.; Carslaw, K. S. Contribution of Feldspar and Marine Organic Aerosols to Global Ice Nucleating Particle Concentrations. *Atmospheric Chem*. Phys. 2017, 17 (5). 10.5194/acp-17-3637-2017.

(27) Šantl-Temkiv, T.; Sikoparija, B.; Maki, T.; Carotenuto, F.; Amato, P.; Yao, M.; Morris, C. E.; Schnell, R.; Jaenicke, R.; Pöhlker, C.; DeMott, P. J.; Hill, T. C. J.; Huffman, J. A. Bioaerosol Field Measurements: Challenges and Perspectives in Outdoor Studies. Aerosol Sci. Technol. 2020, 54 (5), 520–546. 10.1080/02786826.2019.1676395.

(28) Murray, B. J.; O’Sullivan, D.; Atkinson, J. D.; Webb, M. E. Ice Nucleation by Particles Immersed in Supercooled Cloud Droplets. Chem. Soc. Rev. 2012, 41 (19), 6519–6554. 10.1039/c2cs35200a.

(29) Lee, B. U.; Kim, S. H.; Kim, S. S. Hygroscopic Growth of E. Coli and B. Subtilis Bioaerosols. J. Aerosol Sci. 2002, 33 (12), 1721–1723. 10.1016/s0021-8502(02)00114-3.

(30) Sloth Nielsen, L.; Santl-Temkiv, T.; Massling, A.; Finster, K. W.; Bilde, M.; Rosati, B. Hygroscopicity of Airborne Cells of P. Syringae Measured with a Hygroscopicity Tandem Differential Mobility Analyzer. Atmos Chem Phys.

(31) Franc, G. D.; DeMott, P. J. Cloud Activation Characteristics of Airborne Erwinia Carotovora Cells. J. Appl. Meteorol. 2002. 10.1175/1520-0450(1998)037<1293:cacoae>2.0.co;2.

(32) Bauer, H.; Giebl, H.; Hitzenberger, R.; Kasper-Giebl, A.; Reischl, G.; Zibuschka, F.; Puxbaum, H. Airborne Bacteria as Cloud Condensation Nuclei. J. Geophys. Res. Atmospheres 2003, 108 (D21), 1–5. 10.1029/2003jd003545.

(33) Simpson, E. L.; Connolly, P. J.; Mcfiggans, G. Competition for Water Vapour Results in Suppression of Ice Formation in Mixed-Phase Clouds. 2018, 7237–7250.

(34) Kovalev, V. A.; Eichinger, W. E. Elastic Lidar; 2005. 10.1002/0471643173.

(35) Weitkamp, C. Lidarl: Range-Resolved Optical Remote Sensing of the Atmosphere; 2005.

(36) Liou, K. N.; Lahore, H. LASER SENSING OF CLOUD COMPOSITION: A BACKSCATTERED DEPOLARIZATION TECHNIQUE. J. Appl. Meteorol. 1974, 13 (2). 10.1175/1520-0450(1974)013<0257:LSOCCA>2.0.CO;2.

(37) Wang, L.; Stanič, S.; Eichinger, W.; Song, X.; Zavrtanik, M. Development of an Automatic Polarization Raman LiDAR for Aerosol Monitoring over Complex Terrain. Sens. Switz. 2019, 19 (14). 10.3390/s19143186.

(38) Yau, M. K.; Rogers, R. R. A Short Course in Cloud Physics, 3rd edition.; Butterworth-Heinemann: Woburn, Mass., 1989.

(39) Drinovec, L.; Sciare, J.; Stavroulas, I.; Bezantakos, S.; Pikridas, M.; Unga, F.; Savvides, C.; Višić, B.; Remškar, M.; Močnik, G. A New Optical-Based Technique for Real-Time Measurements of Mineral Dust Concentration in PM_10_ Using a Virtual Impactor. Atmospheric Meas. Tech. 2020, 13 (7), 3799–3813. 10.5194/amt-13-3799-2020.

(40) Amin, H.; Marshall, I. P. G.; Bertelsen, R. J.; Wouters, I. M.; Schlünssen, V.; Sigsgaard, T.; Šantl-Temkiv, T. Optimization of Bacterial DNA and Endotoxin Extraction from Settled Airborne Dust. Sci. Total Environ. 2022, 857 (October 2022). 10.1016/j.scitotenv.2022.159455.

(41) Herlemann, D. P. R.; Labrenz, M.; Jürgens, K.; Bertilsson, S.; Waniek, J. J.; Andersson, A. F. Transitions in Bacterial Communities along the 2000 Km Salinity Gradient of the Baltic Sea. ISME J. 2011, 5 (10). 10.1038/ismej.2011.41.

(42) Toju, H.; Tanabe, A. S.; Yamamoto, S.; Sato, H. High-Coverage ITS Primers for the DNA-Based Identification of Ascomycetes and Basidiomycetes in Environmental Samples. PLoS ONE 2012, 7 (7). 10.1371/journal.pone.0040863.

43. Team, R. C. R: A language and environment for statistical computing v. 3.6. 1 (R Foundation for Statistical Computing, Vienna, Austria, 2019). Scientific Reports.

(44) Martin, M. Cutadapt Removes Adapter Sequences from High-Throughput Sequencing Reads. EMBnet.journal 2011. 10.14806/ej.17.1.200.

(45) Callahan, B. J.; McMurdie, P. J.; Rosen, M. J.; Han, A. W.; Johnson, A. J. A.; Holmes, S. P. DADA2: High-Resolution Sample Inference from Illumina Amplicon Data. Nat. Methods 2016, 13 (7), 581–583. 10.1038/nmeth.3869.

(46) Quast, C.; Pruesse, E.; Yilmaz, P.; Gerken, J.; Schweer, T.; Yarza, P.; Peplies, J.; Glöckner, F. O. The SILVA Ribosomal RNA Gene Database Project: Improved Data Processing and Web-Based Tools. Nucleic Acids Res. 2013, 41 (D1). 10.1093/nar/gks1219.

47. UNITE Community. UNITE General FASTA Release for Fungi. Version 18.11.2018, 2018.

(48) Davis, N. M.; Proctor, Di. M.; Holmes, S. P.; Relman, D. A.; Callahan, B. J. Simple Statistical Identification and Removal of Contaminant Sequences in Marker-Gene and Metagenomics Data. Microbiome 2018, 6 (1). 10.1186/s40168-018-0605-2.

(49) McMurdie, P. J.; Holmes, S. Phyloseq: An R Package for Reproducible Interactive Analysis and Graphics of Microbiome Census Data. PLoS ONE 2013, 8 (4). 10.1371/journal.pone.0061217.

(50) Oksanen, J.; Blanchet, F. G.; Friendly, M.; Kindt, R.; Legendre, P.; Mcglinn, D.; Minchin, P. R.; O’Hara, R. B.; Simpson, G. L.; Solymos, P.; Stevens, M. H. H.; Szoecs, E.; Wagner, H. Vegan: Community Ecology Package. R Package Version 2.4-2. Community Ecol. Package 2019, 2.5-6.

(51) Wickham, H. Package ‘ggplot2’: Elegant Graphics for Data Analysis. Springer-Verl. N. Y. 2016.

(52) Wickham, H.; François, R.; Henry, L.; Müller, K. Dplyr: A Grammar of Data Manipulation. R Package Version. Media, 2019.

(53) Wieber, C.; Rosenhøj Jeppesen, M.; Finster, K.; Melvad, C.; Šantl-Temkiv, T. Micro- PINGUIN: Microtiter-Plate-Based Instrument for Ice Nucleation Detection in Gallium with an Infrared Camera. Atmospheric Meas. Tech. 2024, 17 (9), 2707–2719. 10.5194/amt-17-2707-2024.

(54) Vali, G. Quantitative Evaluation of Experimental Results an the Heterogeneous Freezing Nucleation of Supercooled Liquids. J. Atmospheric Sci. 1971, 28 (3), 402–409. 10.1175/1520-0469(1971)028<0402:qeoera>2.0.co;2.

(55) Schwidetzky, R.; Lukas, M.; YazdanYar, A.; Kunert, A. T.; Pöschl, U.; Domke, K. F.; Fröhlich-Nowoisky, J.; Bonn, M.; Koop, T.; Nagata, Y.; Meister, K. Specific Ion– Protein Interactions Influence Bacterial Ice Nucleation. Chem. - Eur. J. 2021, 27 (26), 7402–7407. 10.1002/chem.202004630.

(56) Veselovskii, I.; Goloub, P.; Podvin, T.; Tanre, D.; Ansmann, A.; Korenskiy, M.; Borovoi, A.; Hu, Q.; Whiteman, D. N. Spectral Dependence of Backscattering Coefficient of Mixed Phase Clouds over West Africa Measured with Two-Wavelength Raman Polarization Lidar: Features Attributed to Ice-Crystals Corner Reflection. J. Quant. Spectrosc. Radiat. Transf. 2017, 202. 10.1016/j.jqsrt.2017.07.028.

(57) Korolev, A.; Isaac, G. A. Relative Humidity in Liquid, Mixed-Phase, and Ice Clouds. J. Atmospheric Sci. 2006, 63 (11), 2865–2880. 10.1175/jas3784.1.

(58) Tignat-Perrier, R.; Dommergue, A.; Thollot, A.; Keuschnig, C.; Magand, O.; Vogel, T. M.; Larose, C. Global Airborne Microbial Communities Controlled by Surrounding Landscapes and Wind Conditions. Sci. Rep. 2019, 9 (1). 10.1038/s41598-019-51073-4.

(59) Lee, J.; Song, J.; Lee, D.; Pang, Y. Metal-Enhanced Fluorescence and Excited State Dynamics of Carotenoids in Thin Polymer Films. Sci. Rep. 2019, 9 (1), 1–13. 10.1038/s41598-019-40446-4.

(60) Elleder, M.; Borovanský, J. Autofluorescence of Melanins Induced by Ultraviolet Radiation and near Ultraviolet Light. A Histochemical and Biochemical Study. Histochem. J. 2001, 33 (5), 273–281. 10.1023/A:1017925023408.

(61) Surre, J.; Saint-Ruf, C.; Collin, V.; Orenga, S.; Ramjeet, M.; Matic, I. Strong Increase in the Autofluorescence of Cells Signals Struggle for Survival. Sci. Rep. 2018, 8 (1), 1–14. 10.1038/s41598-018-30623-2.

(62) Tong, Y.; Lighthart, B. Solar Radiation Is Shown to Select for Pigmented Bacteria in the Ambient Outdoor Atmosphere. Photochem. Photobiol. 1997, 65 (1), 103–106. 10.1111/j.1751-1097.1997.tb01884.x.

(63) Amato, P.; Parazols, M.; Sancelme, M.; Laj, P.; Mailhot, G.; Delort, A. M. Microorganisms Isolated from the Water Phase of Tropospheric Clouds at the Puy de Dôme: Major Groups and Growth Abilities at Low Temperatures. In FEMS Microbiology Ecology; 2007. 10.1111/j.1574-6941.2006.00199.x.

(64) Šantl-Temkiv, T.; Finster, K.; Dittmar, T.; Hansen, B. M.; Thyrhaug, R.; Nielsen, N. W.; Karlson, U. G. Hailstones: A Window into the Microbial and Chemical Inventory of a Storm Cloud. PLoS ONE 2013, 8 (1). 10.1371/journal.pone.0053550.

65. Shapiro, Howard M. Practical Flow Cytometry, 4th Edition | Wiley. Wiley.com. https://www.wiley.com/en-us/Practical+Flow+Cytometry%2C+4th+Edition-p-9780471411253 (accessed 2024-07-08).

(66) Stopelli, E.; Conen, F.; Morris, C. E.; Herrmann, E.; Bukowiecki, N.; Alewell, C. Ice Nucleation Active Particles Are Efficiently Removed by Precipitating Clouds. Sci. Rep. 2015, 5, 16433. 10.1038/srep16433.

(67) Stopelli, E.; Conen, F.; Guilbaud, C.; Zopfi, J.; Alewell, C.; Morris, C. E. Ice Nucleators, Bacterial Cells and Pseudomonas Syringae in Precipitation at Jungfraujoch. Biogeosciences 2017, 14 (5), 1189–1196. 10.5194/BG-14-1189-2017.

(68) Amato, P.; Joly, M.; Schaupp, C.; Attard, E.; Möhler, O.; Morris, C. E.; Brunet, Y.; Delort, A.-M. Survival and Ice Nucleation Activity of Bacteria as Aerosols in a Cloud Simulation Chamber. *Atmospheric Chem*. Phys. 2015, 15 (11), 6455–6465. 10.5194/acp-15-6455-2015.

(69) Dusek, U.; Frank, G. P.; Hildebrandt, L.; Curtius, J.; Schneider, J.; Walter, S.; Chand, D.; Drewnick, F.; Hings, S.; Jung, D.; Borrmann, S.; Andreae, M. O. Size Matters More than Chemistry for Cloud-Nucleating Ability of Aerosol Particles. Science 2006, 312 (5778), 1375–1378. 10.1126/science.1125261.

(70) Alsved, M.; Holm, S.; Christiansen, S.; Smidt, M.; Ling, M.; Šantl-temkiv, T. Effect of Aerosolization and Drying on the Viability of Pseudomonas Syringae Cells. Front. Microbiol. 2018, 9 (December), 1–11. 10.3389/fmicb.2018.03086.

(71) Sun, J.; Ariya, P. A. Atmospheric Organic and Bio-Aerosols as Cloud Condensation Nuclei (CCN): A Review. Atmos. Environ. 2006. 10.1016/j.atmosenv.2005.05.052.

(72) Sharma, P. K.; Hanumantha Rao, K. Analysis of Different Approaches for Evaluation of Surface Energy of Microbial Cells by Contact Angle Goniometry. Adv. Colloid Interface Sci. 2002, 98 (3), 341–463. 10.1016/S0001-8686(02)00004-0.

73. Hashmi, I.; Bindschedler, S.; Junier, P. Chapter 18 - Firmicutes. In Beneficial Microbes in Agro-Ecology; Amaresan, N., Senthil Kumar, M., Annapurna, K., Kumar, K., Sankaranarayanan, A., Eds.; Academic Press, 2020; pp 363–396. 10.1016/B978-0-12-823414-3.00018-6.

(74) Zboralski, A.; Filion, M. Pseudomonas Spp. Can Help Plants Face Climate Change. Front. Microbiol. 2023, 14. 10.3389/fmicb.2023.1198131.

(75) Frohlich-Nowoisky, J.; Pickersgill, D. A.; Despres, V. R.; Poschl, U. High Diversity of Fungi in Air Particulate Matter. Proc. Natl. Acad. Sci. 2009, 106 (31), 12814–12819. 10.1073/pnas.0811003106.

(76) Fröhlich-Nowoisky, J.; Burrows, S. M.; Xie, Z.; Engling, G.; Solomon, P. A.; Fraser, M. P.; Mayol-Bracero, O. L.; Artaxo, P.; Begerow, D.; Conrad, R.; Andreae, M. O.; Després, V. R.; Pöschl, U. Biogeography in the Air: Fungal Diversity over Land and Oceans. Biogeosciences 2012, 9, 1125–1136. 10.5194/bg-9-1125-2012.

(77) Archer, S. D. J.; Lee, K. C.; Caruso, T.; Alcami, A.; Araya, J. G.; Cary, S. C.; Cowan, D. A.; Etchebehere, C.; Gantsetseg, B.; Gomez-Silva, B.; Hartery, S.; Hogg, I. D.; Kansour, M. K.; Lawrence, T.; Lee, C. K.; Lee, P. K. H.; Leopold, M.; Leung, M. H. Y.; Maki, T.; McKay, C. P.; Al Mailem, D. M.; Ramond, J.-B.; Rastrojo, A.; Šantl-Temkiv, T.; Sun, H. J.; Tong, X.; Vandenbrink, B.; Warren-Rhodes, K. A.; Pointing, S. B. Contribution of Soil Bacteria to the Atmosphere across Biomes. Sci. Total Environ. 2023, 871, 162137. 10.1016/j.scitotenv.2023.162137.

(78) Chase, A. B.; Arevalo, P.; Polz, M. F.; Berlemont, R.; Martiny, J. B. H. Evidence for Ecological Flexibility in the Cosmopolitan Genus Curtobacterium. Front. Microbiol. 2016, 7 (NOV), 1–11. 10.3389/fmicb.2016.01874.

(79) Sampaio, J. P. Cystofilobasidium Oberwinkler & Bandoni (1983). The Yeasts 2011, 3 (1983), 1423–1432. 10.1016/B978-0-444-52149-1.00111-7.

(80) Vasebi, Y.; Mechan Llontop, M. E.; Hanlon, R.; Schmale III, D. G.; Schnell, R.; Vinatzer, B. A. Comprehensive Characterization of an Aspen (*Populus Tremuloides*) Leaf Litter Sample That Maintained Ice Nucleation Activity for 48 Years. Biogeosciences 2019, 16 (8), 1675–1683. 10.5194/bg-16-1675-2019.

(81) Agarkova, I. V.; Lambrecht, P. A.; Vidaver, A. K.; Harveson, R. M. Genetic Diversity among Curtobacterium Flaccumfaciens Pv. Flaccumfaciens Populations in the American High Plains. Can. J. Microbiol. 2012, 58 (6). 10.1139/W2012-052.

(82) Trutko, S. M.; Dorofeeva, L. V.; Evtushenko, L. I.; Ostrovskii, D. N.; Hintz, M.; Wiesner, J.; Jomaa, H.; Baskunov, B. P.; Akimenko, V. K. Isoprenoid Pigments in Representatives of the Family Microbacteriaceae. Mikrobiologiya 2005, 74 (3), 335–341.

(83) Vanek, M.; Mravec, F.; Szotkowski, M.; Byrtusova, D.; Haronikova, A.; Certik, M.; Shapaval, V.; Marova, I. Fluorescence Lifetime Imaging of Red Yeast Cystofilobasidium Capitatum during Growth. EuroBiotech J. 2018, 2 (2), 114–120. 10.2478/ebtj-2018-0015.

84. Abusada, G. M. Studies on the Mechanism of Protection by Carotenoids. PhD Thesis 1992.

(85) Furutani, H.; Dall’osto, M.; Roberts, G. C.; Prather, K. A. Assessment of the Relative Importance of Atmospheric Aging on CCN Activity Derived from Field Observations. Atmos. Environ. 2008, 42 (13), 3130–3142. 10.1016/j.atmosenv.2007.09.024.

(86) Xia, Y.; Conen, F.; Alewell, C. Total Bacterial Number Concentration in Free Tropospheric Air above the Alps. Aerobiologia 2013, 29 (1), 153–159. 10.1007/s10453-012-9259-x.

(87) Bowers, R. M.; Lauber, C. L.; Wiedinmyer, C.; Hamady, M.; Hallar, A. G.; Fall, R.; Knight, R.; Fierer, N. Characterization of Airborne Microbial Communities at a High-Elevation Site and Their Potential to Act as Atmospheric Ice Nuclei. Appl. Environ. Microbiol. 2009, 75 (15), 5121–5130. 10.1128/AEM.00447-09.

(88) Joly, M.; Amato, P.; Deguillaume, L.; Monier, M.; Hoose, C.; Delort, A. M. Quantification of Ice Nuclei Active at near 0 °c Temperatures in Low-Altitude Clouds at the Puy de Dôme Atmospheric Station. *Atmospheric Chem*. Phys. 2014, 14 (15). 10.5194/acp-14-8185-2014.

(89) Petters, M. D.; Wright, T. P. Revisiting Ice Nucleation from Precipitation Samples. Geophys. Res. Lett. 2015, 42 (20), 8758–8766. 10.1002/2015GL065733.

(90) Dedekind, Z.; Proske, U.; Ferrachat, S.; Lohmann, U.; Neubauer, D. Simulating the Seeder–Feeder Impacts on Cloud Ice and Precipitation over the Alps. *Atmospheric Chem*. Phys. 2024, 24 (9), 5389–5404. 10.5194/acp-24-5389-2024.

(91) Minder, J. R.; Mote, P. W.; Lundquist, J. D. Surface Temperature Lapse Rates over Complex Terrain: Lessons from the Cascade Mountains. J. Geophys. Res. Atmospheres 2010, 115 (D14). 10.1029/2009JD013493.

(92) Zufall, M. J.; Davidson, C. I.; Caffrey, P. F.; Ondov, J. M. Airborne Concentrations and Dry Deposition Fluxes of Particulate Species to Surrogate Surfaces Deployed in Southern Lake Michigan. Environ. Sci. Technol. 1998, 32 (11). 10.1021/es9706458.

(93) Caffrey, P. F.; Ondov, J. M.; Zufall, M. J.; Davidson, C. I. Determination of Size-Dependent Dry Particle Deposition Velocities with Multiple Intrinsic Elemental Tracers. Environ. Sci. Technol. 1998, 32 (11). 10.1021/es970644f.

(94) Williams, R. M. A Model for the Dry Deposition of Particles to Natural Water Surfaces. Atmospheric Environ. 1967 1982, 16 (8). 10.1016/0004-6981(82)90464-4

(95) Qi, J.; Li, P.; Li, X.; Feng, L.; Zhang, M. Estimation of Dry Deposition Fluxes of Particulate Species to the Water Surface in the Qingdao Area, Using a Model and Surrogate Surfaces. Atmos. Environ. 2005, 39 (11), 2081–2088. 10.1016/j.atmosenv.2004.12.017.

