## Supplemental Fig 1-18 for "The limited capacity of bioaerosols to serve as cloud-condensation nuclei may restrict their potential to initiate ice formation in mixed-phase clouds"

**Supplementary material**

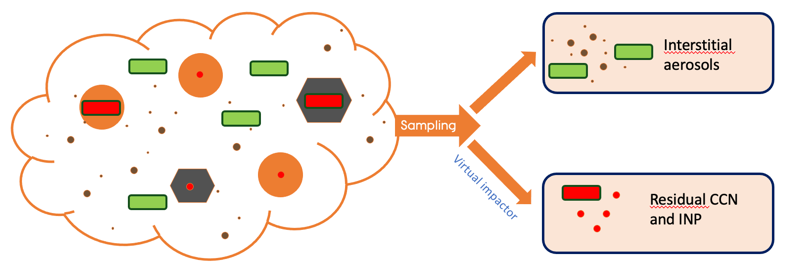

SI-Figure 1: Aerosols from the two cloud phases were collected separately as interstitial and residual aerosols using a virtual impactor.

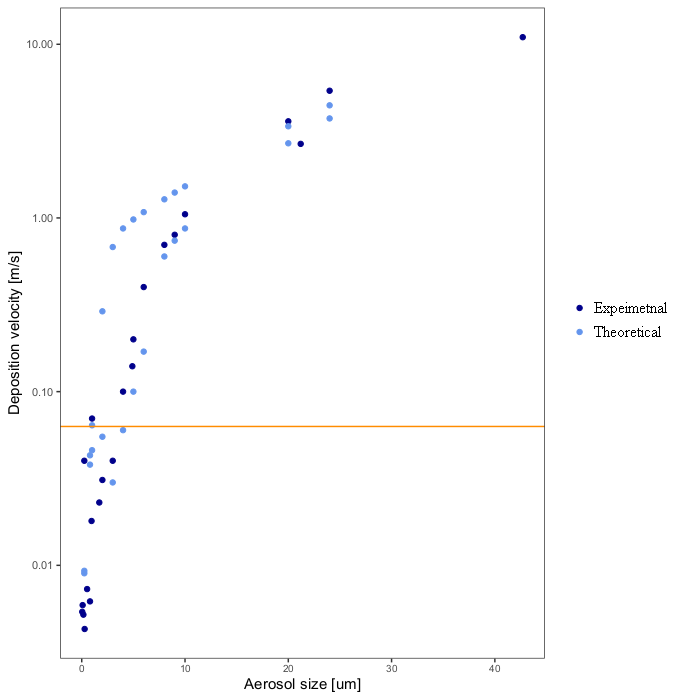

SI-Figure 2: Deposition velocity as a function of aerosol size. The experimental ^92,93^ and theoretical ^94,95^ values were all obtained from ^95^. The orange horizontal line indicates the threshold deposition velocity, below which the particles will get collected with the assumed BioSampler flow of 12 lpm.

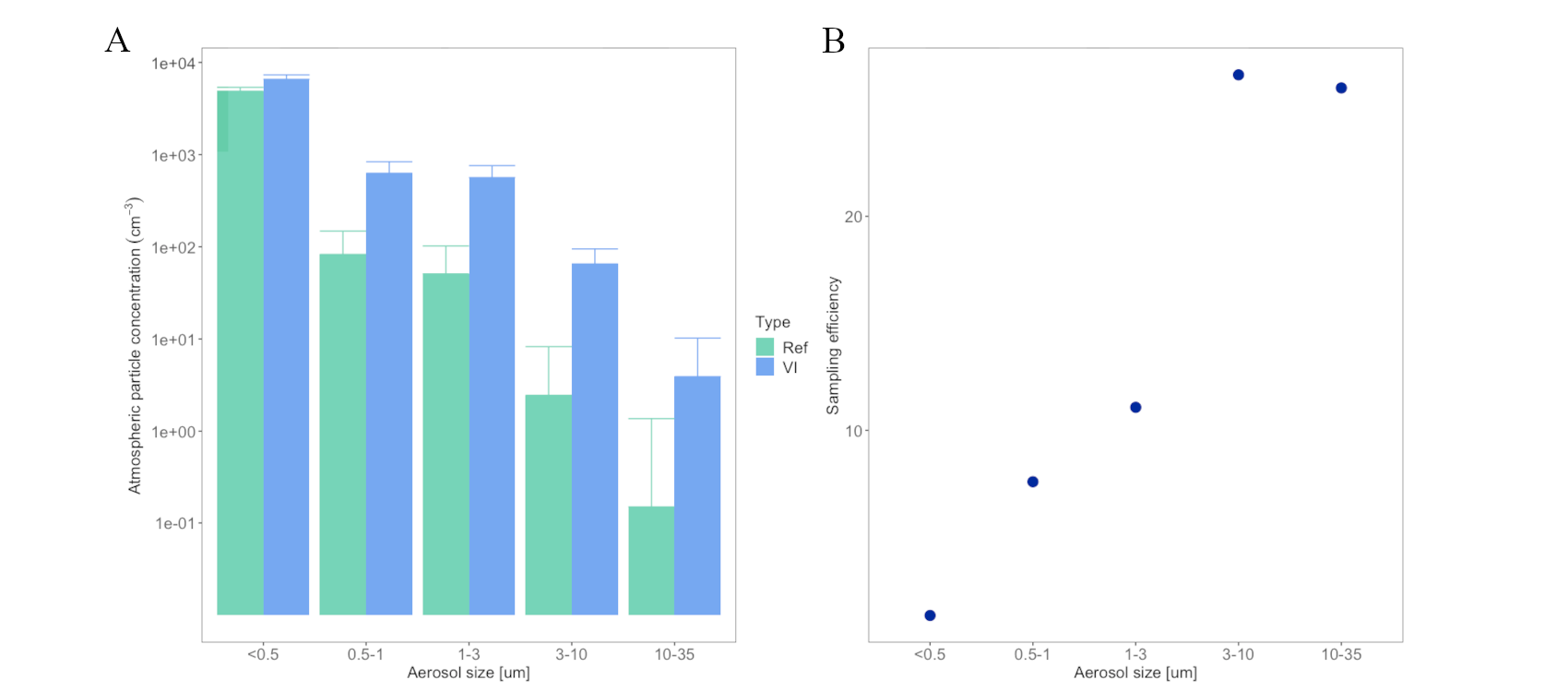

SI-Figure 3: (A) Atmospheric particle concentration as determined by the optical particle size spectrometer (OPSS) for different aerosol size ranges as determined for total indoor laboratory air (Ref) and as determined when the OPSS was coupled with the virtual impactor (VI) inlet using a total-to-minor flow ratio of 6.3-7.3. The measurements for the Ref and the VI were carried out for 20 min, respectively. (B) The concentration efficiency as a function of atmospheric particle size.

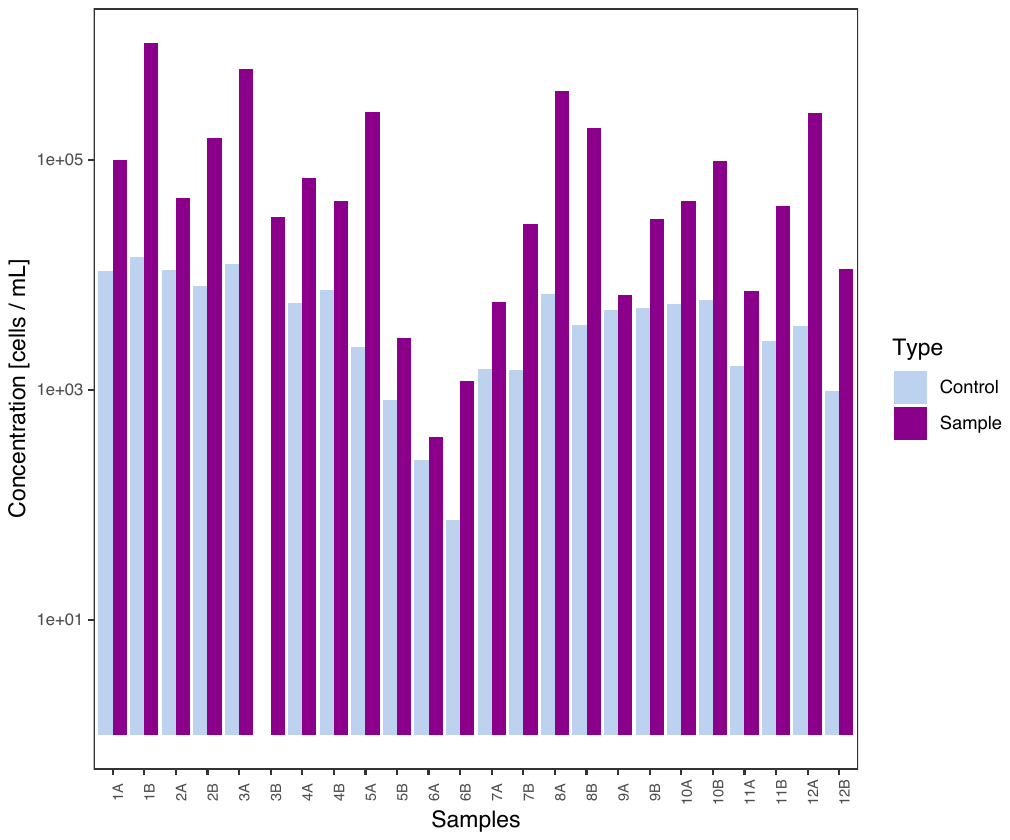

SI-Figure 4: The concentration of cells per volume of liquid in samples and corresponding controls as determined by flow cytometry.

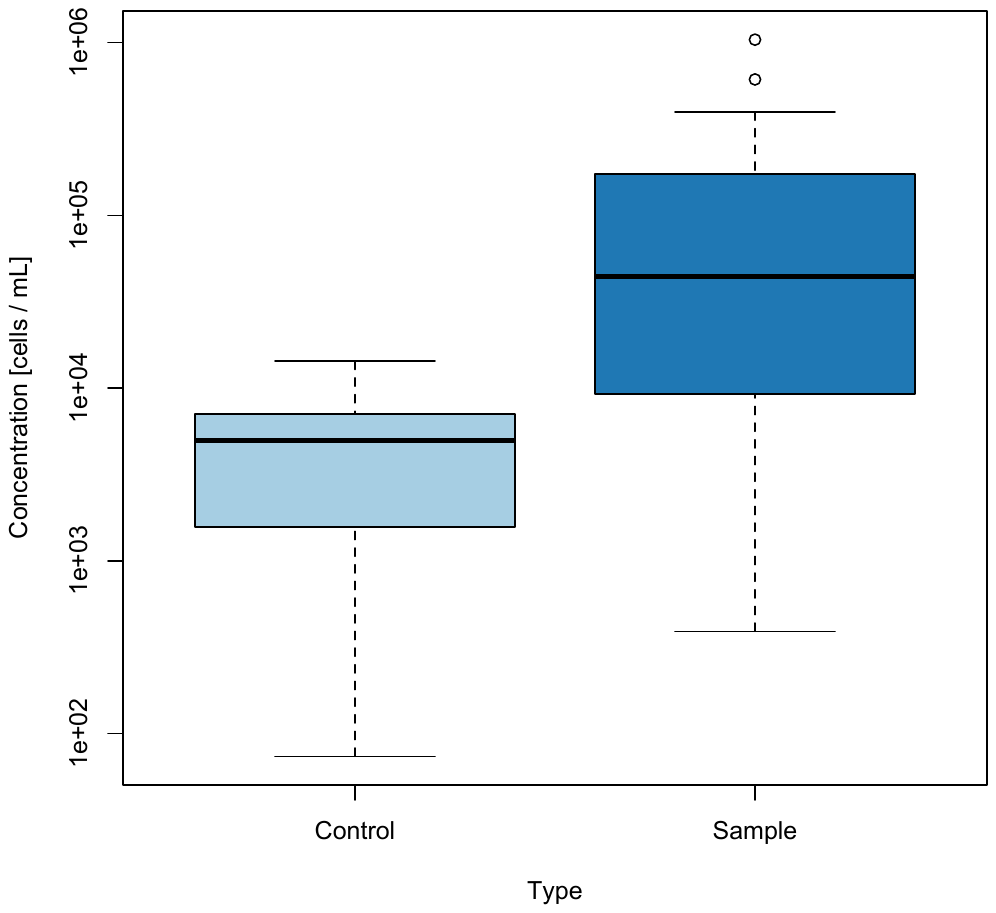

SI-Figure 5: A bar plot showing the average concentration of cells per volume of liquid in samples and corresponding controls as determined by flow cytometry.

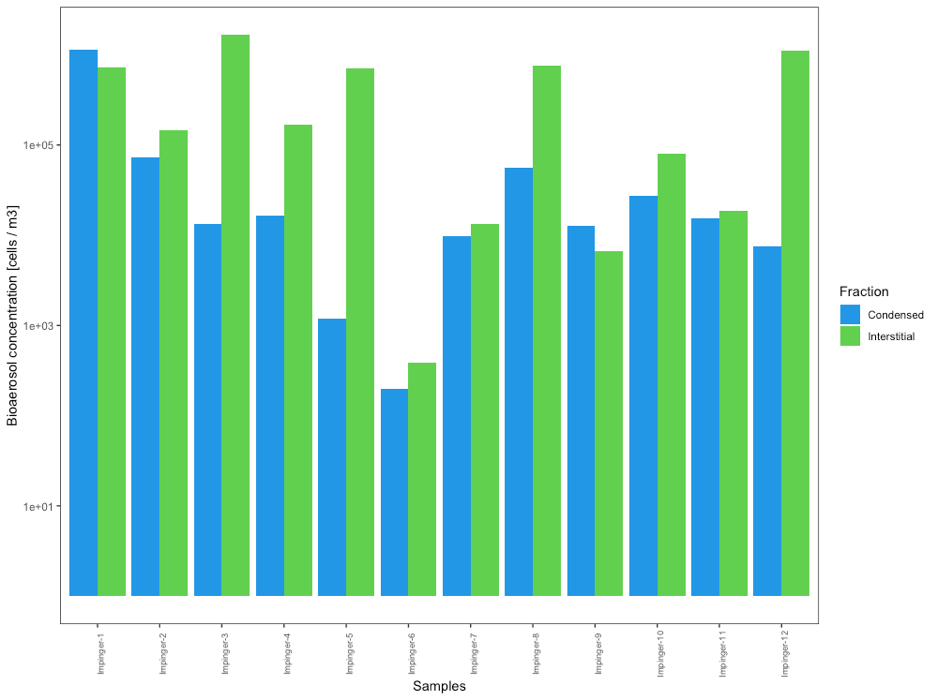

SI-Figure 6: Bioaerosol concentration per volume of air for the individual sampling points and the condensed as well as the interstitial phases of the clouds.

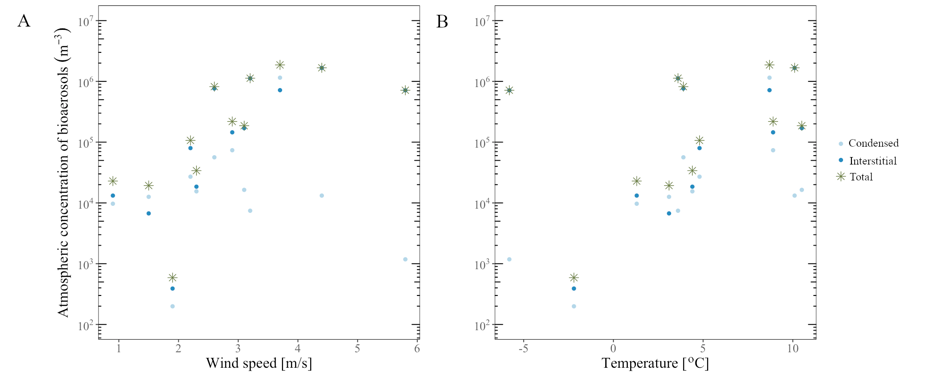

SI-Figure 7: The relationship between the atmospheric concentration of bioaerosols and (A) wind speed; (B) temperature.

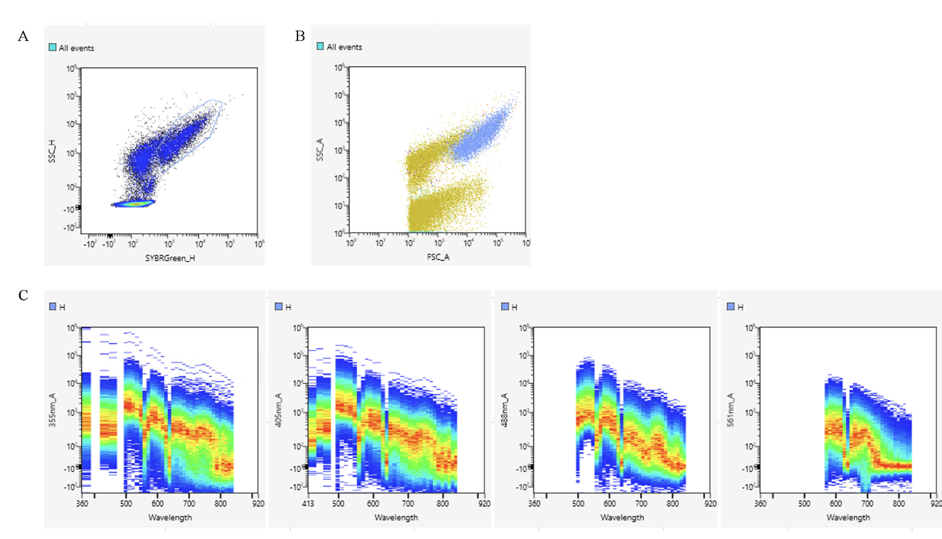

SI-Figure 8: Dot-plots and spectral properties showing the predominant autofluorescent population as determined by flow cytometry using the ID7000 (Sony Biotechnology) spectral flow cytometer on unstained samples. (A) Side scatter is shown as a function of fluorescence in the B2 channel (SYBRGreen_H axis). The blue gate encircles the autofluorescent population (HF cells). (B) Side scatter is shown as a function of the forward scatter. The autofluorescent population is shown in light blue, while the rest of the events are in green. (C) The spectral signature as obtained with each laser for the autofluorescent population.

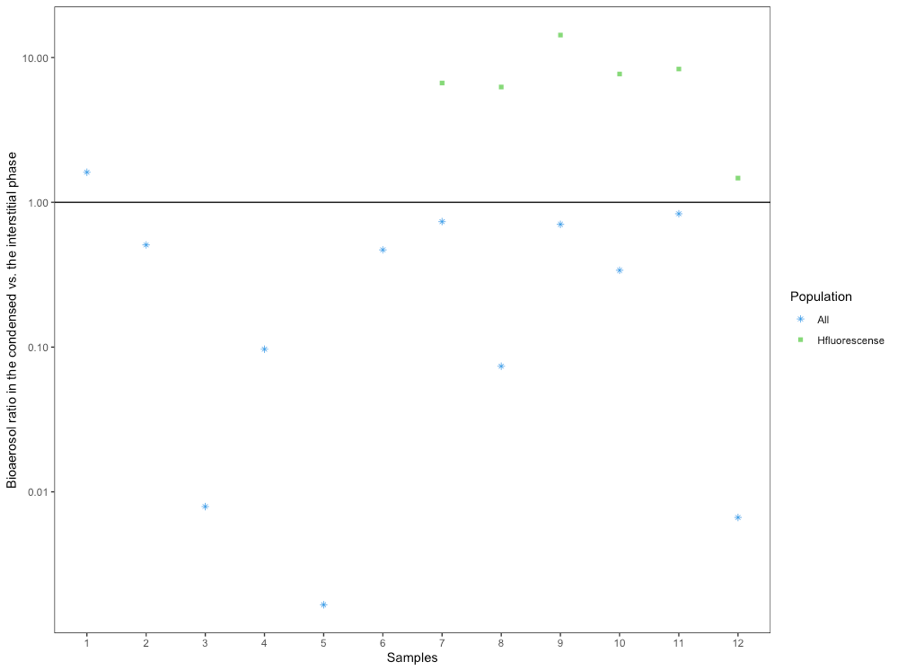

SI-Figure 9: The ratio between activated and interstitial bioaerosols in individual samples showing both all bioaerosols (“All”) and bioaerosols within the autofluorescent population (“Hfluorescence”) as quantified by flow cytometry.

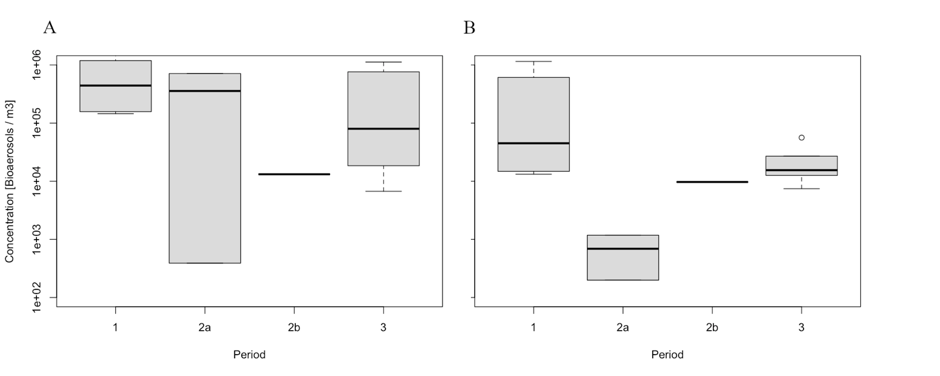

SI-Figure 10: The concentration of bioaerosols in the (A) interstitial and (B) condensed phase during the three campaigns. During the second campaign 2a refers to the concentrations during cloud events and 2b refers to the concentrations when there was no cloud.

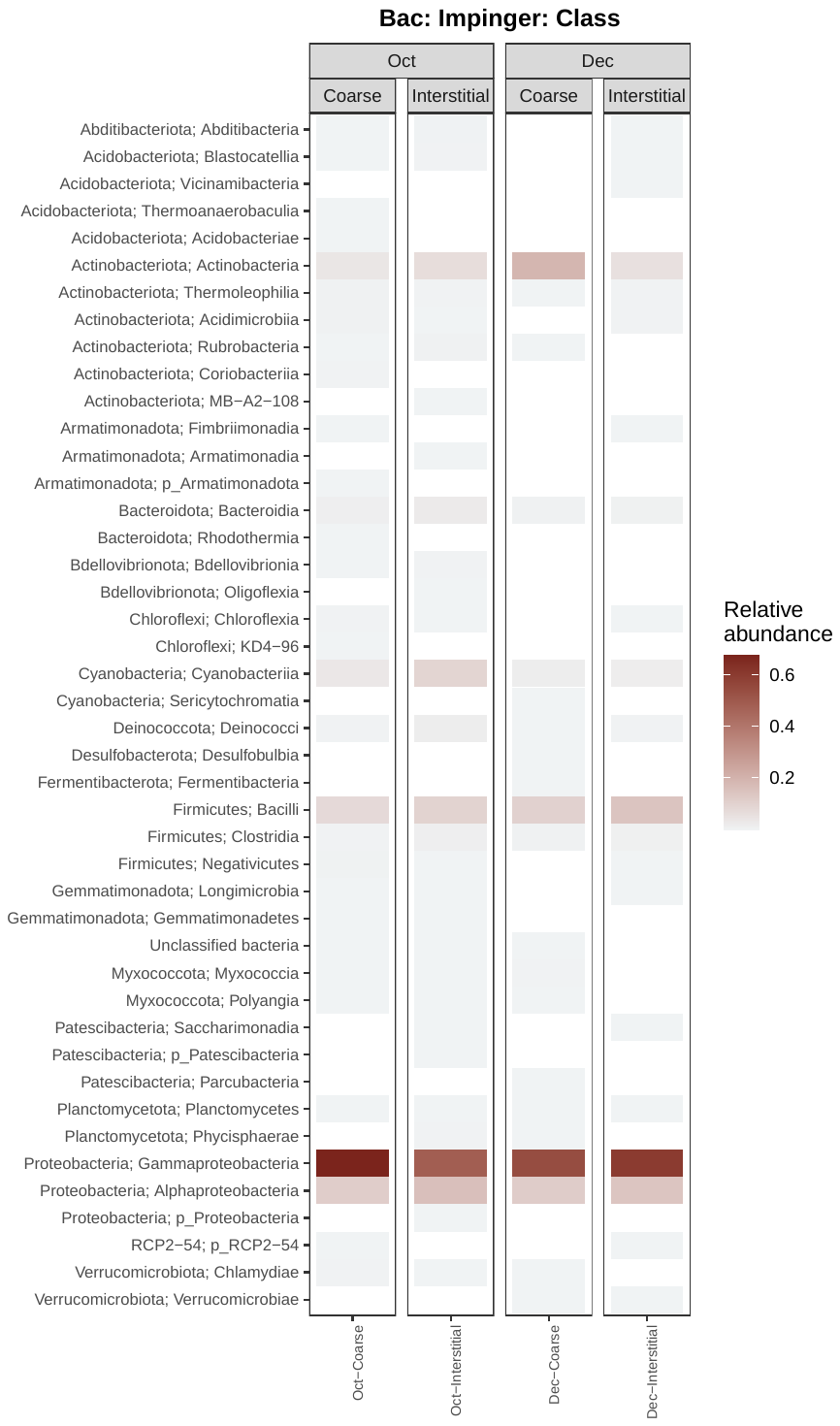

SI-Figure 11: A heatmap showing the average relative abundance of major bacterial classes for the condensed and interstitial phases in October and December.

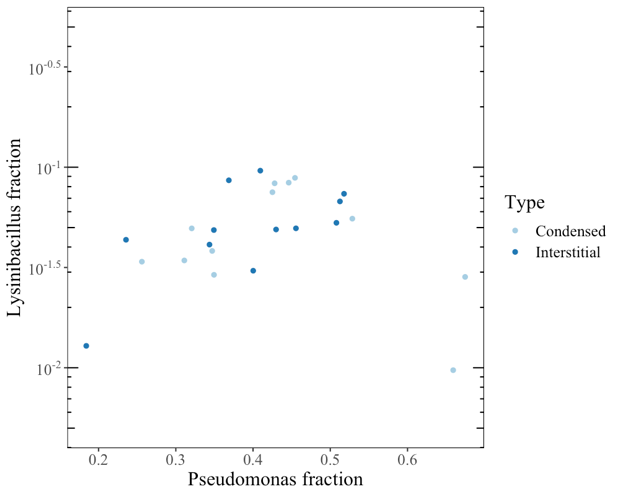

SI-Figure 12: Ratio of *Lysinibacillus* sp. as a function of the ration of *Pseudomonas* sp. in condensed and interstitial phases of the clouds.

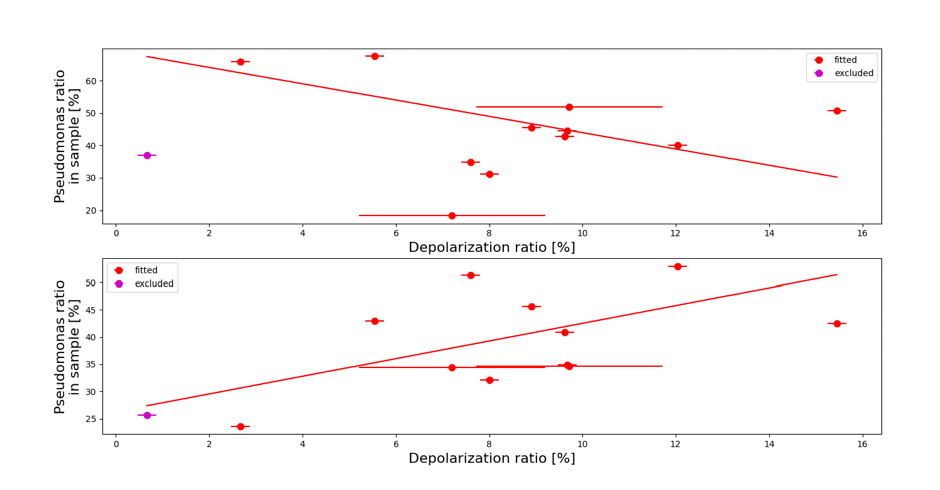

SI-Figure 13: Ratio of *Pseudomonas* sp., the predominant bacteria genus, in condensed and interstitial phase with respect to depolarization. Linear fit shows a decreasing trend (−2.52 ± 1.14) in the condensed phase and an increasing trend (1.62 ± 0.74) in the interstitial phase. The data point in magenta corresponds to the case when there was no cloud at measurement height and was not used in calculation of the correlation.

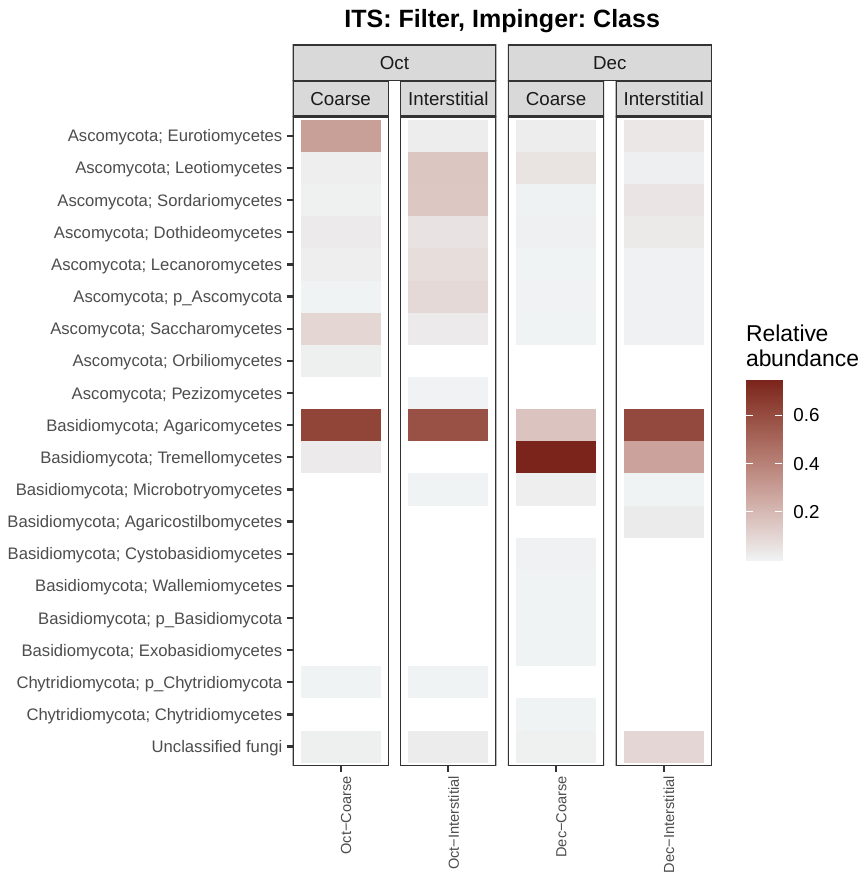

SI-Figure 14: A heatmap showing the average relative abundance of major fungal classes for the condensed and interstitial phases in October and December.

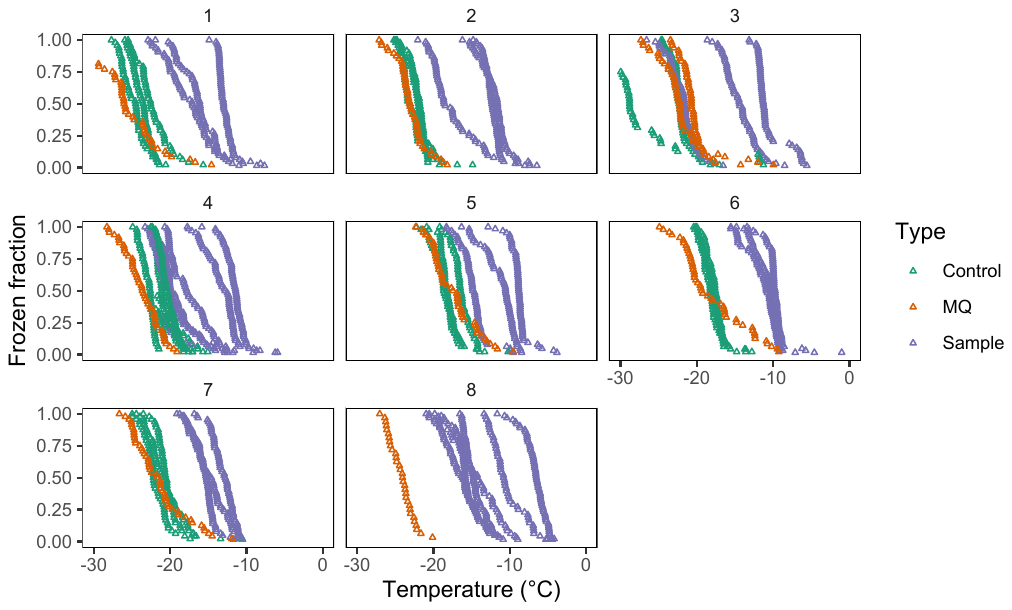

SI-Figure 15: The frozen fraction as a function of the temperature obtained for each ice-nucleation measurement performed.

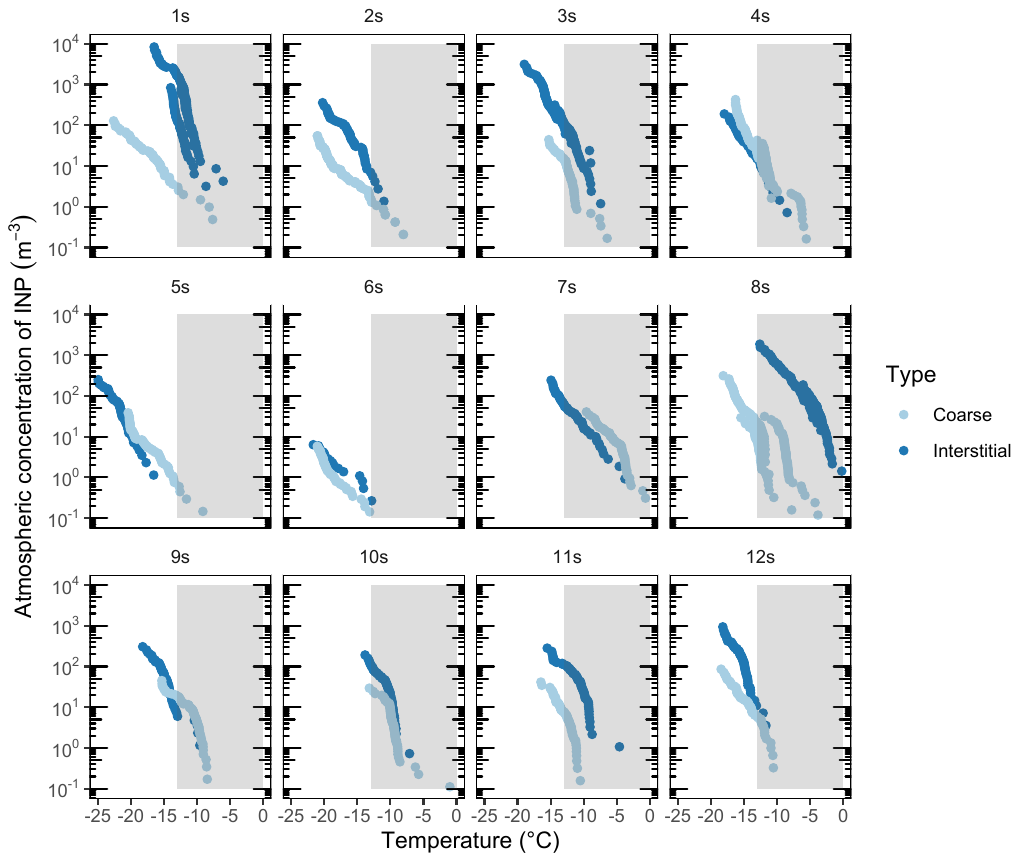

SI-Figure 16: The ice-nucleation spectra for each cloud event showing the comparison of the condensed and interstitial cloud phases. The shaded area denotes biogenic INP.

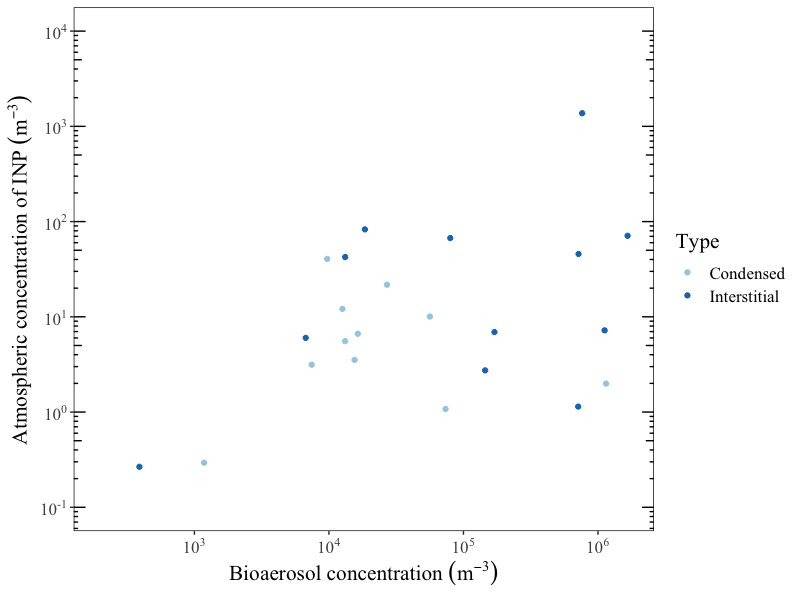

SI-Figure 17: The relationship between the atmospheric concentration of INP_-12_ and bioaerosol concentration.

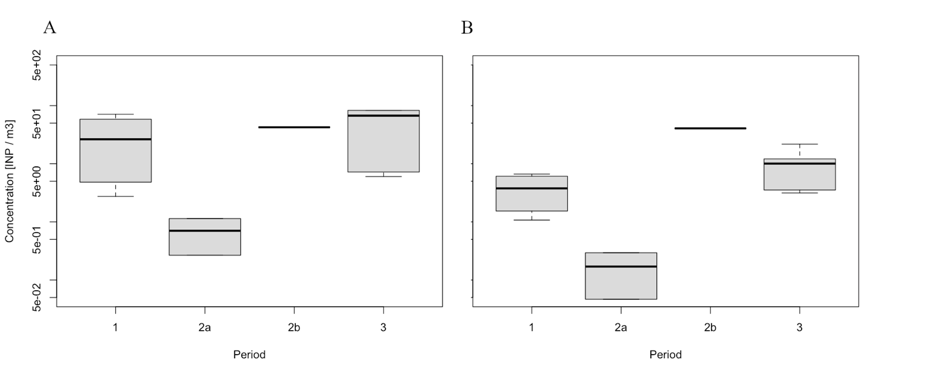

SI-Figure 18: The concentration of INP-12 in the (A) interstitial and (B) condensed phase during the three campaigns. During the second campaign 2a refers to the concentrations during cloud events and 2b refers to the concentrations when there was no cloud.
